# Neural surprise in somatosensory Bayesian learning

**DOI:** 10.1101/2020.06.18.158915

**Authors:** Sam Gijsen, Miro Grundei, Robert T. Lange, Dirk Ostwald, Felix Blankenburg

**Author notes:** Corresponding authors (SG), (MG). These authors contributed equally to this work (Sam Gijsen, Miro Grundei).

## Abstract

Tracking statistical regularities of the environment is important for shaping human behavior and perception. Evidence suggests that the brain learns environmental dependencies using Bayesian principles. However, much remains unknown about the employed algorithms, for somesthesis in particular. Here, we describe the cortical dynamics of the somatosensory learning system to investigate both the form of the generative model as well as its neural surprise signatures. Specifically, we recorded EEG data from 40 participants subjected to a somatosensory roving-stimulus paradigm and performed single-trial modeling across peri-stimulus time in both sensor and source space. Our Bayesian model selection procedure indicates that evoked potentials are best described by a non-hierarchical learning model that tracks transitions between observations using leaky integration. From around 70ms post-stimulus onset, secondary somatosensory cortices are found to represent confidence-corrected surprise as a measure of model inadequacy. Primary somatosensory cortex is found to encode Bayesian surprise, reflecting model updating, from around 140ms. As such, this dissociation indicates that early surprise signals may control subsequent model update rates. In sum, our findings support the hypothesis that early somatosensory processing reflects Bayesian perceptual learning and contribute to an understanding of its precise mechanisms.

**Author summary:** Our environment features statistical regularities, such as a drop of rain predicting imminent rainfall. Despite the importance for behavior and survival, much remains unknown about how these dependencies are learned, particularly for somatosensation. As surprise signalling about novel observations indicates a mismatch between one’s beliefs and the world, it has been hypothesized that surprise computation plays an important role in perceptual learning. By analyzing EEG data from human participants receiving sequences of tactile stimulation, we compare different formulations of surprise and investigate the employed underlying learning model. Our results indicate that the brain estimates transitions between observations. Furthermore, we identified different signatures of surprise computation and thereby provide a dissociation of the neural correlates of belief inadequacy and belief updating. Specifically, early surprise responses from around 70ms were found to signal the need for changes to the model, with encoding of its subsequent updating occurring from around 140ms. These results provide insights into how somatosensory surprise signals may contribute to the learning of environmental statistics.

## Introduction

The world is governed by statistical regularities, such that a single drop of rain on the skin might predict further tactile sensations through imminent rainfall. The learning of such probabilistic dependencies facilitates adaptive behaviour and ultimately survival. Building on ideas tracing back to Helmholtz [1], it has been suggested that the brain employs an internal generative model of the environment which generates predictions of future sensory input. More recent accounts of perception and perceptual learning, including predictive coding [2, 3] and the free energy principle [4], propose that these models are continuously updated in light of new sensory evidence using Bayesian inference. Under such a view, the generative model is composed of a likelihood function of sensory input given external causes and a prior probability distribution over causes [4, 5]. Perception is interpreted as the computation of a posterior distribution over causes of sensory input and model parameters, while perceptual learning is seen as the updating of the prior distribution based on the computed posterior [6]. Such a description of Bayesian perceptual learning has been successfully used to explain aspects of learning in the auditory [7–9], visual [10–12], as well as somatosensory domain [13].

To investigate the underlying neuronal dynamics of perceptual inference, predictions formed by the brain can be probed by violating statistical regularities. Widely researched neurobiological markers of regularity violation include EEG components such as the auditory mismatch negativity (aMMN) and the P300 in response to deviant stimuli following regularity inducing standard stimuli. As an alternative to the oddball paradigm typically used to elicit such mismatch responses (MMRs) [14], the roving-stimulus paradigm features stimulus sequences that alternate between different trains of repeated identical stimuli [15]. Expectations are built up across a train of stimuli of variable length and are subsequently violated by alternating to a different stimulus train. The paradigm thereby allows for the study of MMRs based on the sequence history and independently of the physical stimulus properties. Analogues to the aMMN have also been reported for vision [16] and somatosensation (sMMN). The sMMN was first reported by Kekoni et al. [17] and has since been shown in response to deviant stimuli with different properties, including spatial location [18–26], vibrotactile frequency [17, 27–29], and stimulus duration [30, 31]. Increasing evidence has been reported for an account of the MMN as a reflection of Bayesian perceptual learning processes for the auditory [32, 33, 8], visual [16, 12], and to a lesser extent the somatosensory domain [13]. However, the precise mechanisms remain unknown, as it is unclear whether the MMN reflects the signaling of the inadequacy of the current beliefs (prediction error) or their adjustment, due to the lack of direct comparisons between these competing accounts.

In the context of probabilistic inference, the signalling of a mismatch between predicted and observed sensory input may be formally described using computational quantities of surprise [6, 34]. By adopting the vocabulary introduced by Faraji et al. [35] surprise can be grouped into two classes: puzzlement and enlightenment surprise. Puzzlement surprise refers to the initial realization of a mismatch between the world and an internal model. Predictive surprise (PS) captures this concept based on the measure of information as introduced by Shannon [36]. Specifically, PS considers the frequency of events such that the occurrence of a rare event (i.e. associated with low probability of occurring) is more informative and results in greater surprise. It was recently proposed to consider the commitment to a generative model in addition to the amount of information conveyed by an observation. For example, in order for the percept of a drop of rain on the skin to be surprising, commitment to a belief about a clear sky may be necessary. This idea of puzzlement surprise is quantified by the measure of confidence-corrected surprise (CS) [35]. The concept of enlightenment surprise, on the other hand, directly relates to the size of the update of the world model that may follow initial puzzlement. Bayesian surprise (BS) [37, 38] captures this notion by quantifying the degree to which an observer adapts their internal generative model in order to accommodate novel observations [37, 38]. Both predictive surprise [9] and Bayesian surprise [13] have been successfully applied to the full time-window of peri-stimulus EEG data to model neural surprise signals. However, the majority of studies have focused on P300 amplitudes, with applications of both predictive surprise [39–42] and Bayesian surprise [43, 40, 44]. Earlier EEG signals have received less attention, although the MMN was reported to reflect PS [42]. Furthermore, due to the close relationship between model updating and prediction violation, only few studies have attempted to dissociate their signals. Although the use of different surprise functions in principle allows for a direct comparison of the computations potentially underlying EEG mismatch responses, such studies remain scarce. Previous research either focused on their spatial identification using fMRI [11, 45–47] or temporally specific, late EEG components [40]. Finally, to the best of our knowledge, only one recent pre-print study compared all three prominent surprise functions in a reanalysis of existing data, reporting PS to be better decoded across the entire post stimulus time-window [48].

Despite the successful account of perceptual learning using Bayesian approaches, the frame-work is broad and much remains unclear about the nature of MMRs, their description as surprise signals, and the underlying generative models that give rise to them. This is especially the case for the somatosensory modality, though evidence has been reported for the encoding of Bayesian surprise using the roving paradigm [13]. The current study expands on this work by recording EEG responses to a roving paradigm formulated as a generative model with discrete hidden states. We explore different mismatch responses, including the somatosensory analogue to the MMN, independent of the physical properties of stimuli. Using single-trial modeling, we systematically investigate the structure of the generative model employed by the brain. Having established the most likely probabilistic model, we provide a spatiotemporal description of its different surprise signatures in electrode and source space. As direct comparisons are scarce, we thus contribute by dissecting the dynamics of multiple aspects of Bayesian computation utilized for somatosensory learning across peri-stimulus time by incorporating them into one hierarchical analysis.

## Materials and Methods

### Experimental Design

#### Participants

44 healthy volunteers (18-38 years old, mean age: 26, 28 females, all right-handed) participated for monetary compensation of 10 Euro per hour or an equivalent in course credit. The study was approved by the local ethics committee of the Freie Universität Berlin and written informed consent was obtained from all subjects prior to the experiment.

#### Experimental Procedure

In order to study somatosensory mismatch responses and model them as single-trial surprise signals, we used an oddball-like roving-stimulus paradigm [15]. Stimuli were applied in consecutive trains of alternating stimuli based on a probabilistic model (see below) with an inter-stimulus interval of 750ms (see Fig 1). Trains of stimuli consisted of two possible stimulation intensities. The first and last stimulus in a train were labeled as a deviant and standard, respectively. Thus, as opposed to a classic oddball design, the roving paradigm allows for both stimulus types to function as a standard or deviant.

**Fig 1.**
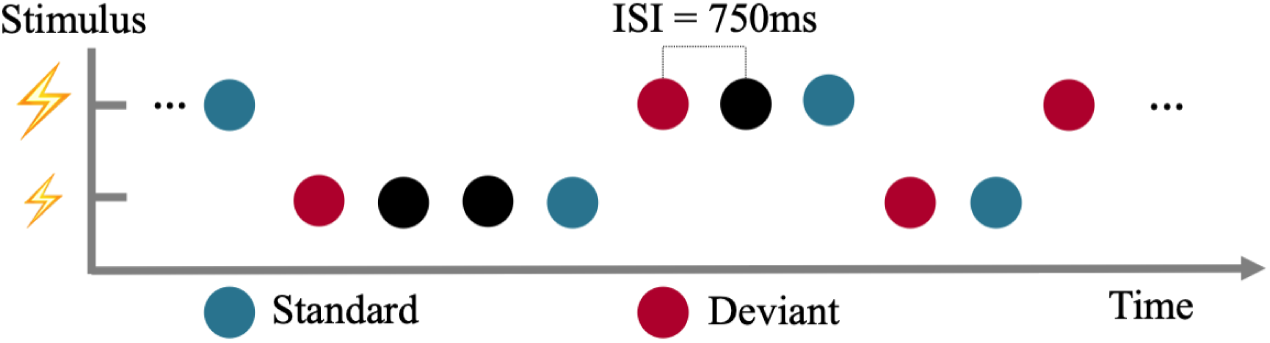
Presentation of experimental stimuli using a roving-like oddball paradigm. Stimuli with two different intensities are presented. Their role as standard or deviant depends on their respective position within the presentation sequence.

Adhesive electrodes (GVB-geliMED GmbH, Bad Segeberg, Germany) were attached to the wrist through which the electrical stimuli with a 0.2ms duration were administered. In order to account for interpersonal differences in sensory thresholds, the two intensity levels were determined on a subject basis. The low intensity level (mean 5.05mA±1.88) was set in proximity to the detection threshold yet so that stimuli were clearly perceivable. The high intensity level (mean 7.16mA±1.73) was determined for each subject to be easily distinguishable from the low intensity level, yet remaining non-painful and below the motor threshold. The catch stimulus (described below) featured a threefold repetition of the 0.2ms stimulus at an interval of 50ms and was presented at either the low or high intensity level with equal probability.

Following familiarization with the electrical stimulation, 800 stimuli were administered in each of 5 experimental runs à 10 minutes. To ensure the subjects maintained attention on the electrical stimulation, they were instructed to count the number of catch trials (targets). In order to make the task non-trivial, the probability of the occurrence of a catch stimulus was set to either 0.01, 0.015, 0.02, 0.025, or 0.03, corresponding to a range of 3-32 trials per run. A subject received a stimulus sequence corresponding to each catch trial probability only once, with the order randomized between subjects. Following an experimental run, subjects indicated their counted number of catch trials and received feedback in the form of the correct amount.

### EEG data collection and preprocessing

Data were collected using a 64-channel active electrode system (ActiveTwo, BioSemi, Amsterdam, Netherlands) at a sampling rate of 2048Hz, with head electrodes placed in accordance to the extended 10-20 system. Individual electrode positions were digitalized and recorded using an electrode positioning system (zebris Medical GmbH, Isny, Germany) with respect to three fiducial markers placed on the subject’s face; left and right preauricular points and the nasion. This approach aided subsequent source reconstruction analyses.

Preprocessing was performed using SPM12 (Wellcome Trust Centre for Neuroimaging, Institute for Neurology, University College London, London, UK) and in-house scripts. First, the data were referenced against the average reference, high-pass filtered (0.01Hz), and downsampled to 512Hz. Consequently, eye-blinks were corrected using a topological confound approach [49] and epoched using a peri-stimulus time interval of −100 to 600ms. All trials were then visually inspected and removed in case any significant artefacts were deemed to be present. The EEG data of four subjects were found to contain excessive noise due to hardware issues, resulting in their omission from further analyses and leaving 40 subjects. Finally, a low-pass filter was applied (45Hz). Grand mean somatosensory evoked potentials (SEPs) were calculated for deviant stimuli (‘deviants’) and for the standard stimuli directly preceding a deviant to balance the number of trials (‘standards’). The preproccesed EEG data was baseline corrected with respect to the pre-stimulus interval of −100 to −5 ms. For the GLM analyses, each trial of the electrode data was subsequently linearly interpolated into a 32×32 plane for each timepoint, resulting in a 32×32×308 image per trial. To allow for the use of random field theory to control for family-wise errors, the images were smoothed with a 12 by 12 mm full-width half-maximum (FWHM) Gaussian kernel. Catch trials were omitted for both the ERP and single-trial analyses.

### Generation of stimuli sequences

A property of generative models that is highly relevant for learning in dynamic environments is the manner by which they may adapt their estimated statistics in the face of environmental changes. By incorporating occasional switches between sets of sequence statistics, we aimed to compare generative models that embody different mechanisms of adapting to such change-points. Specifically, the sequential presentation of the stimuli originated from a partially observable probabilistic model for which the hidden state evolved according to a Markov chain (Fig 2) with 3 states *s*. The state transition (*p*(*s*_*t*_|*s*_*t*−1_)) and emission probabilities *p*(*o*_*t*_|*o*_*t*−1_, *o*_*t*−2_, *s*_*t*_) of the observations *o* are listed in Table 1. One of the states was observable as it was guaranteed to emit a catch trial, while the other two states were latent, resembling fast and slow switching regimes. As the latter was specified with higher transition probabilities associated with repeating observations (*p*(0|00) and *p*(0|01)) it thus produced longer stimulus trains on average. For every run, the sequence was initialized by starting either in the slow or fast switching regime with equal probability (*p*(*s*_1_) = {0.5, 0.5, 0}, with catch probability being 0) and likewise producing a high or low stimulus with equal probability (*p*(*o*_1_|*s*_1_) = {0.5, 0.5}).

**Table 1.**
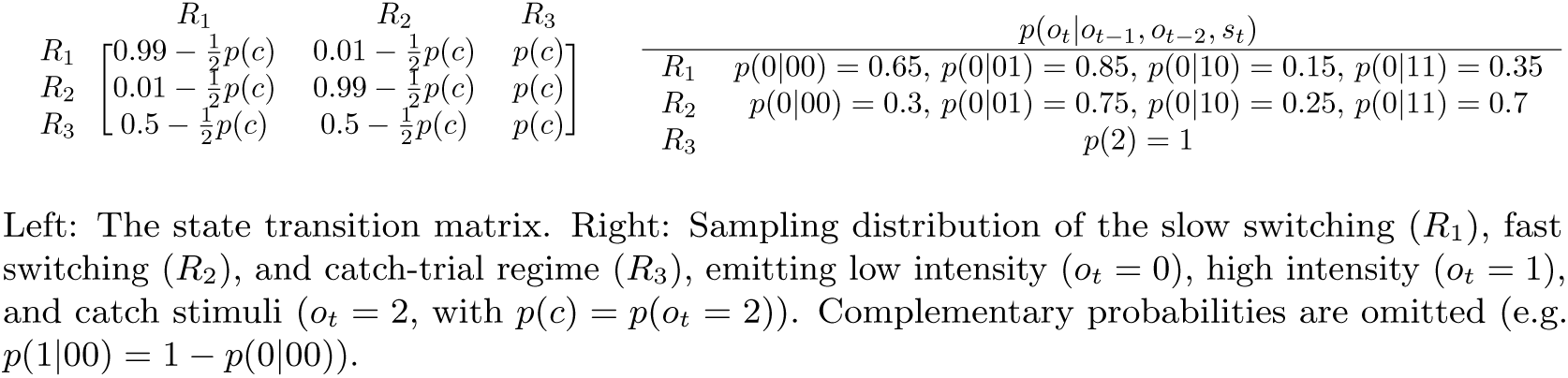
Data-generating process.

| | $R_1$ | $R_2$ | $R_3$ | $p(o_t\vert o_{t-1}, o_{t-2}, s_t)$ | |
| --- | --- | --- | --- | --- | --- |
| $R_1$ | $0.99 - \frac{1}{2}p(c)$ | $0.01 - \frac{1}{2}p(c)$ | $p(c)$ | $R_1$ | $p(0\vert 00) = 0.65, p(0\vert 01) = 0.85, p(0\vert 10) = 0.15, p(0\vert 11) = 0.35$ |
| $R_2$ | $0.01 - \frac{1}{2}p(c)$ | $0.99 - \frac{1}{2}p(c)$ | $p(c)$ | $R_2$ | $p(0\vert 00) = 0.3, p(0\vert 01) = 0.75, p(0\vert 10) = 0.25, p(0\vert 11) = 0.7$ |
| $R_3$ | $0.5 - \frac{1}{2}p(c)$ | $0.5 - \frac{1}{2}p(c)$ | $p(c)$ | $R_3$ | $p(2) = 1$ |

Left: The state transition matrix. Right: Sampling distribution of the slow switching ( $R_1$ ), fast switching ( $R_2$ ), and catch-trial regime ( $R_3$ ), emitting low intensity ( $o_t = 0$ ), high intensity ( $o_t = 1$ ), and catch stimuli ( $o_t = 2$ , with $p(c) = p(o_t = 2)$ ). Complementary probabilities are omitted (e.g. $p(1|00) = 1 - p(0|00)$ ).

**Fig 2.**
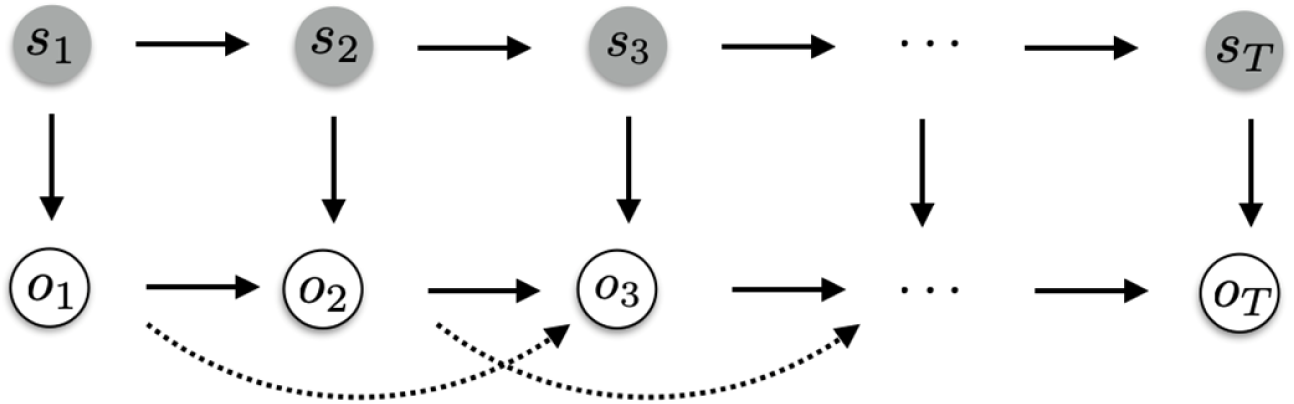
Graphical model of data-generating process. Upper row depicts the evolution of states *s*_*t*_ over time according to a Markov chain. The states emit observations *o*_*t*_ (lower row), which themselves feature second order dependencies on the observation level.

### Event-related potentials

To investigate the event-related response to the experimental conditions on the EEG data, the statistical design was implemented with the general linear model using SPM12. On the first level, the single-trial data of each participant was subject to a multiple regression approach with regressors for each experimental variable: stimulus type (standard/deviant), stimulation regime (fast/slow switching regime), train length ([2, 3, 4, 5, >6] stimuli) and nuisance regressors accounting for experimental blocks (1-5). The restricted maximum likelihood estimation implemented in SPM12 yielded *β*-parameter estimates for each model regressor over (scalp-)space and time which were further analysed at the group level. The second level consisted of a mass-univariate multiple regression analysis of the individual *β* scalp-time images with a design matrix specifying regressors for stimulus type and regime as well as parametric regressors for train length and block and an additional subject factor. The condition contrasts were then computed by weighted summation of the group level regressors’ *β* estimates. To control for multiple comparisons, the scalp-time images were corrected with SPM’s random field theory-based family wise error correction (FWE) [50]. The corresponding *β*-parameter estimates of the significant peaks of the GLM were investigated further with linear regression to test for a linear relationship with train lengths.

### Distributed source localization

In order to establish the somatosensory system as the driving dipolar generator of the EEG signals prior to 200ms, we followed a two-stage source reconstruction analysis consisting of a distributed and an equivalent current dipole (ECD) approach. While we report and model later EEG components in sensor-space, we refrained from source localizing these, as they most likely originate from a more distributed network of multiple sources [51, 52]. Furthermore, the somatosensory system has been shown to be involved in mismatch processing in the time window prior to 200ms [18, 19, 23, 30, 53, 26].

The distributed source reconstruction algorithm as implemented in SPM12 was used to determine the sources of the ERP’s on a subject level. Specifically, subject-specific forward models were created using a 8196 vertex template cortical mesh which was co-registered with the electrode positions using the three aforementioned fiducial markers. SPM12’s BEM EEG head model was used to construct the forward model’s lead field. The multiple sparse priors under group constraints were implemented for the subject-specific source estimates [54, 55]. These were subsequently analyzed at the group level using one-sample t-tests. The yielded statistical parametric maps were thresholded at the peak level with p<0.05 after FWE correction. The anatomical correspondence of the MNI coordinates of the cluster peaks were verified via cytoarchitectonic references using the SPM Anatomy toolbox. Details of the distributed source reconstruction can be reviewed in the results section.

### Equivalent current dipole fitting & source projection

The results of the distributed source reconstruction were subsequently used to fit ECDs to the grand average ERP data using the variational Bayes ECD fitting algorithm implemented in SPM12. The MNI coordinates resulting from the distributed source reconstruction served as informed location priors with variance of 10mm^2^ to optimize the location and orientation of the dipoles for a time-window around the peak of each component of interest (shown in the results section). For the primary somatosensory cortex (S1), two individual dipoles were fit to the time windows of the N20 and P50 components, respectively, to differentiate two sources of early somatosensory processing. Furthermore, a symmetrical dipolar source was fit to the peak of the N140 component of the evoked response with an informed prior around the secondary somatosensory cortex. Subsequently, the single trial EEG data of each subject was projected with the ECD lead fields onto the 4 sources using SPM12, which enabled model selection analyses in source-space.

### Trial-by-trial modeling of sensor- and source-space EEG data

#### Sequential Bayesian learner models for categorical data

To compare Bayesian learners in terms of their generative models and surprise signals, we specified various probabilistic models which generate the regressors ultimately fitted to the EEG data. Capitalizing on the occasional changes to the sequence statistics included in the experimental stimulus generating model, we assess two approaches to latent state inference. Specifically, a conjugate Dirichlet-Categorical (DC) model as well as a Hidden Markov Model (HMM) were used for modeling categorical data. The DC model is non-hierarchical and does not feature any explicit detection of the regime-switches. However, it is able to adapt its estimated statistics to account for sequence change-points by favoring recent observations over those in the past, akin to a progressive “forgetting” or leaky integration. The model assumes a real-valued, static hidden state *s*_*t*_ that is shared across time for each observation emission:

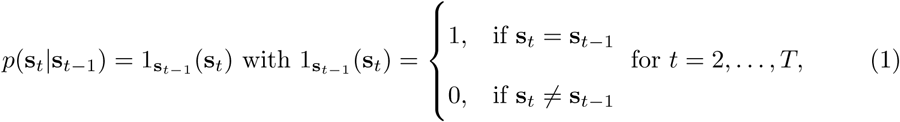

where the probability of the model to be in state **s**_*t*_ is 1 if it equals the previous state and 0 otherwise.

In contrast, the HMM is a hierarchical model for which *s*_*t*_ is a discrete variable and assumed to follow a first order Markov Chain, mimicking the data generation process. As such, it contains additional assumptions about the task structure, which allows for flexible adaptation following a regime-switch by performing inference over a set of discrete hidden states *K* (*s*_*t*_ ∈ {1, …, *K*}). The transition dynamics are given by the row-stochastic matrix **A** ∈ ℝ^*K*×*K*^ with *a*_*ij*_ ≥ 0 and 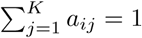:

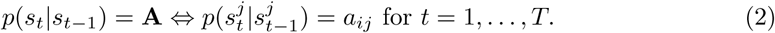

Within our two model classes, we differentiate between four probabilistic models. Here, the aim is to investigate which sequence statistics are estimated by the generative model. In the case of Stimulus Probability (SP) inference, the model does not capture any Markov dependence: *o*_*t*_ solely depends on *s*_*t*_. Alternation Probability (AP) inference captures a limited form of first-order Markov dependency, by estimating the probability of the event of altering observations *d*_*t*_ given the hidden state *s*_*t*_ and the previous observation *o*_*t*−1_, where 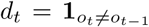 takes on the value 1 if the current observation *o*_*t*_ differs from *o*_*t*−1_. With Transition Probability (TP_1_) inference, the model accounts for full first-order Markov dependence and estimates separate alternation probabilities depending on *o*_*t*−1_ and *s*_*t*_, i.e. *p*(*o*_*t*_|*o*_*t*−1_, *s*_*t*_). Finally, TP_1_ inference may be extended (TP_2_) to also depend on *o*_*t*−2_, and by estimating *p*(*o*_*t*_|*s*_*t*_, *o*_*t*−1_, *o*_*t*−2_) it most closely resembles the structure underlying the data generation.

### Dirichlet-Categorical model

The Dirichlet-Categorical model is part of the Bayesian conjugate pairs. It models the likelihood of the observations using the Cate-gorical distribution with {1, …, *M*} different possible realizations per sample *y*_*t*_. Given the probability vector **s** = {*s*_1_, …, *s*_*M*_} defined on the *M* − 1 dimensional simplex *S*_*M*−1_ with *s*_*i*_ > 0 and 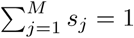, the probability mass function of an event is given by

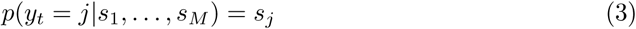

Furthermore, the prior distribution over the hidden state **s** is given by the Dirichlet distribution which is parametrized by the probability vector *α* = {*α*_1_, …, *α*_*M*_}:

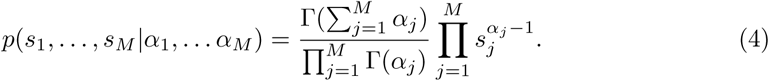

Hence, we have a Dirichlet prior with *s*_1_, …, *s*_*M*_ ∼ *Dir*(*α*_1_, …, *α*_*M*_) and a Categorical likelihood with *y* ∼ *Cat*(*s*_1_, …, *s*_*M*_). Given a sequence of observations *y*_1_, …, *y*_*t*_ the model then combines the likelihood evidence with prior beliefs in order refine posterior estimates over the latent variable space (derivations of enumerated formulas may be found in the supplementary material):

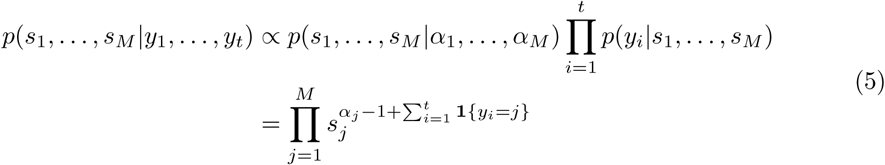

Since the Dirichlet prior and Categorical likelihood pair follow the concept of conjugacy, given an initial 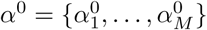 (set as a hyperparameter) the filtering distribution can be computed:

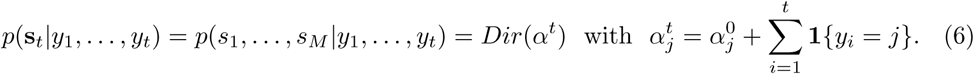

Likewise, one can easily obtain the posterior predictive distribution (needed to compute the predictive surprise readout) by integrating over the space of latent states:

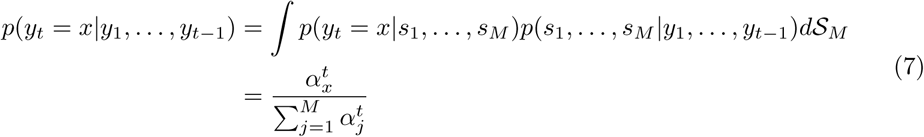

We can evaluate the likelihood of a specific sequence of events which can be used to iteratively compute the posterior:

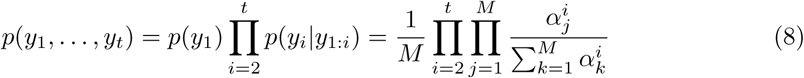

For the evaluation of the posterior distributions, we differentiate between three inference types which track different statistics of the incoming sequence as described above:

1. The stimulus probability (SP) model: *y*_*t*_ = *o*_*t*_ for *t* = 1, …, *T*
2. The alternation probability (AP) model: *y*_*t*_ = *d*_*t*_ for *t* = 2, …, *T*
3. The transition probability model (TP_1_ & TP_2_): *y*_*t*_ = *o*_*t*_ for *t* = 1, …, *T* with a set of hidden parameters **s**_**1**_^(*i*)^ for each transition from *o*_*t*−1_ = *i* and **s**_**2**_^(*j*)^ for each transition from *o*_*t*−2_ = *j* respectively

Despite a static latent state representation, the DC model may account for hidden dynamics by incorporating an exponential memory-decay parameter *τ* ∈ [0, 1] which discounts observations the further in the past they occurred. Functioning as an exponential forgetting mechanism, it allows for the specification of different timescales of observation integration.

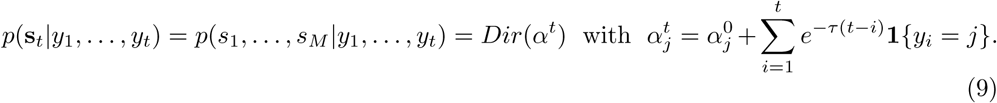

### Hidden Markov Model

While the Dirichlet-Categorical model provides a simple yet expressive conjugate Bayesian model for which analytical posterior expressions exist, it is limited in the functionality of the latent state *s* due to its interpretation as the discrete distribution over categories. Hidden Markov Models (HMMs), on the other hand, are able to capture the dynamics of the hidden state with the transition probabilities of a Markov Chain (MC). Given the hidden state at time *t*, the categorical observation *o*_*t*_ is sampled according to the stochastic matrix **B** ∈ ℝ^*M* ×*K*^, containing the emission probabilities, *p*(*o*_*t*_|*s*_*t*_). The evolution of the discrete hidden state according to a MC, *p*(*s*_*t*_|*s*_*t*−1_), is described by the stochastic matrix **A** ∈ ℝ^*K*×*K*^. The initial hidden state *p*(*s*_1_) is sampled according to the distribution vector *π* ∈ ℝ^*K*^. **A, B** are both row stochastic, hence 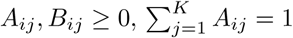 and 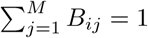. The graphical model described by the HMM setup is thereby specified as depicted in Fig 4.

**Fig 3.**
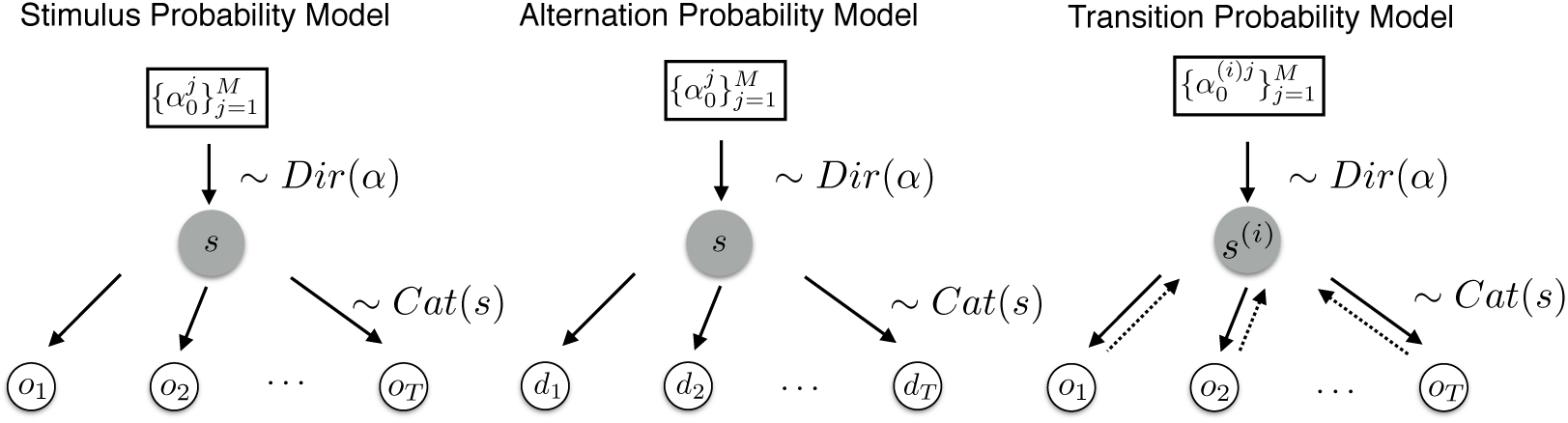
Dirichlet-Categorical model as a graphical model. Left: The stimulus probability model which tracks the hidden state vector determining the sampling process of the raw observations. Middle: The transition probability model which infers the hidden state distribution based on alternations of the observations. Right: The transition probability model which assumes a different data-generating process based on the previous observations. Hence, it infers *M* sets of probability vectors *α*^*i*^.

**Fig 4.**
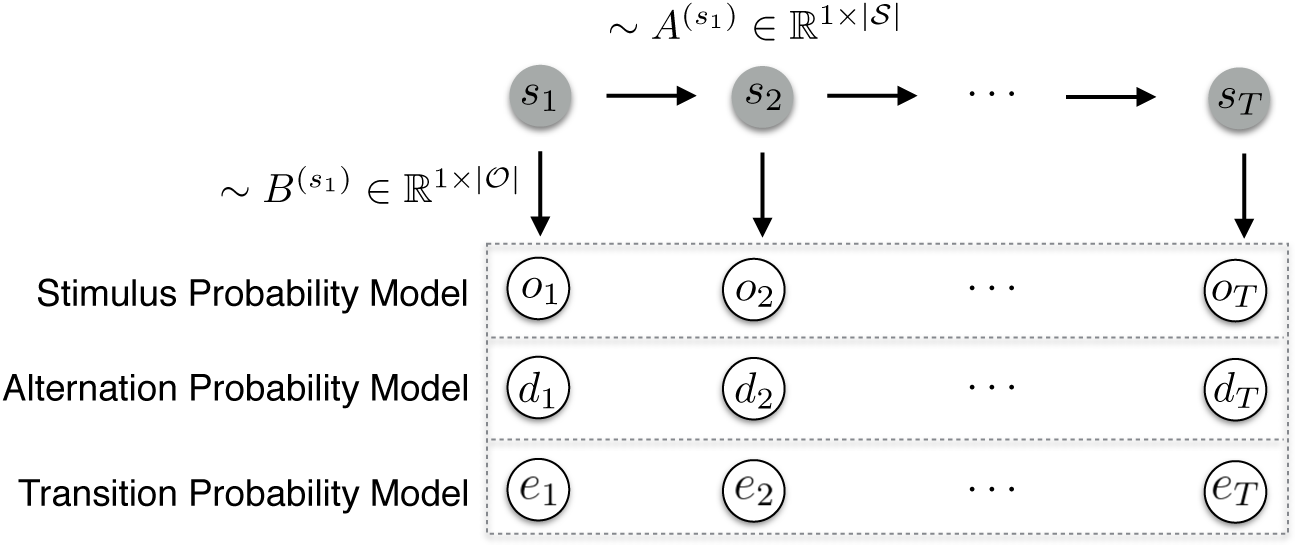
Hidden Markov Model as a graphical model. Upper row depicts the evolution of states *s*_*t*_ according to the transition matrix 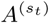. The states emit observational data (dotted rectangle) according to the probabilities specified in stochastic matrix 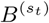 which depends on the type of inference. The stimulus probability model infers the emission probabilities associated with the raw observations *o*_*t*_. The alternation probability model tracks the alternations of observations with 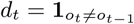. The transition probability model assumes a data-generating process based on previous observations, with *e*_*t*_ coding for the transitions between observations.

Classically, the parameters of this latent variable are inferred using the Expectation-Maximisation (EM) algorithm. Therefore, and in order to derive the factorisation of the joint likelihood *p*(*o*_1:*t*_, *s*_1:*t*_), the backward and forward probabilities are used in conjunction with the Baum-Welch algorithm in order to perform the inference procedure (see S1 Appendix).

### HMM Implementation

The aim of the HMM was to approximate the data generation process more closely by using a model capable of learning the regimes over time and performing latent state inference at each timestep. To this end, prior knowledge was used in its specification by fixing the state transition matrix close to its true values (*p*(*s*_*t*_ = *s*_*t*−1_) = 0.99). The rare catch trials were removed from the data prior to fitting the HMM and thus their accompanying third regime was omitted, resulting in a two-state HMM. Given that an HMM estimates emission probabilities of the form *p*(*o*_*t*_|*s*_*t*_) and thus does not capture any additional explicit dependency on previous observations, the input vector of observations was transformed prior to fitting the models. For AP and TP inference this equated to re-coding the observation *o*_*t*_ to reflect the specific event that occurred. Specifically, for the AP model the input sequence was 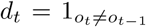, while for TP_1_ and TP_2_ a vector of events was used corresponding to the four possible transitions from *o*_*t*−1_ or eight transitions from *o*_*t*−2_ respectively. Thus, the HMM estimates two vectors of emission probabilities corresponding to these events. Despite this deviation of the fitted models from the underlying data generation process, the AP and TP models reliably captured R_1_ and R_2_ to their capability, with TP_2_ retrieving the true, but unknown underlying emission probabilities (see S1 Fig). As expected, SP inference was agnostic to the regimes, while AP and TP inference allowed for the tracking of the latent state over time (S2 Fig). An example of the filtering posterior may be found in Fig 5.

**Fig 5.**
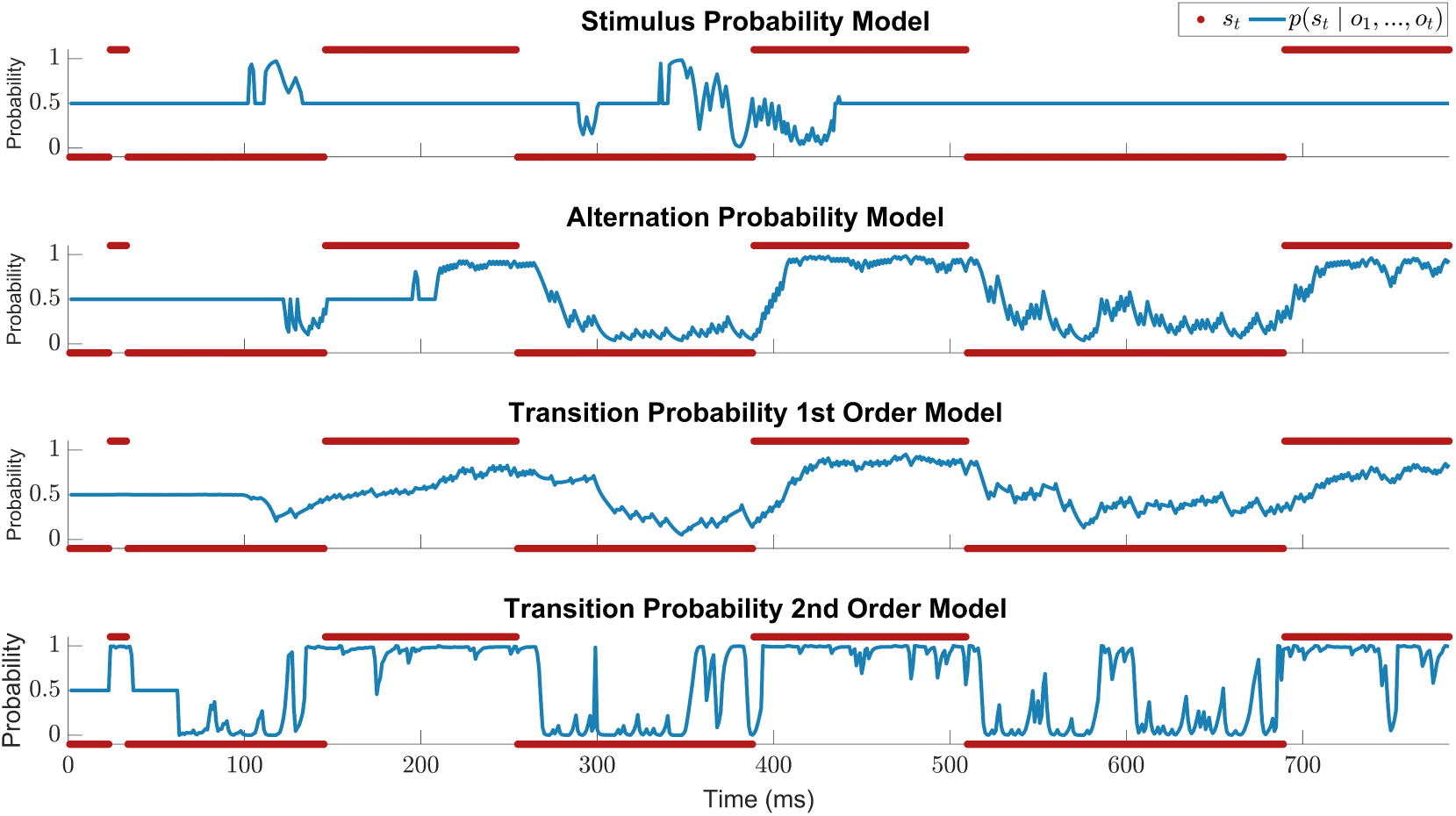
Posterior probabilities of the HMM. Comparison of the posterior *p*(*s*_*t*_ |*o*_1_, …, *o*_*t*_) of the different HMM inference models for an example sequence. The true, but unknown regimes of the data generation process are plotted in red.

### Surprise readouts

For each of the probabilistic models described above, three different surprise functions were implemented, forming the predictors for the EEG data: predictive surprise *PS*(*y*_*t*_), Bayesian surprise *BS*(*y*_*t*_), and confidence-corrected surprise *CS*(*y*_*t*_). These may be interpreted as read-out functions of the generative model, signalling a mismatch between the world and the internal model.

The predictive surprise is defined as the negative logarithm of the posterior predictive distribution *p*(*y*_*t*_|*s*_*t*_):

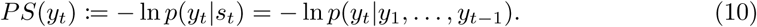

A posterior that assigns little probability to an event *y*_*t*_ will cause high (unit-less) predictive surprise and as such is a measure of puzzlement surprise. The Bayesian surprise, on the other hand, quantifies enlightenment surprise and is defined as the Kullback-Leibler (KL) divergence between the posterior pre- and post-update:

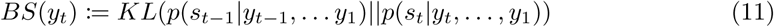

Confidence-corrected surprise is an extended definition of puzzlement surprise which additionally considers the commitment of the generative model as it is scaled by the negative entropy of the prior distribution. It is defined as the KL divergence between the prior and posterior distribution of a naive observer, corresponding to an agent with a flat prior 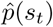 (i.e. all outcomes are equally likely) which observed *y*_*t*_:

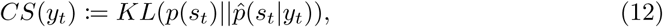

For the DC model, the flat prior 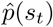 can be written as *Dir*(*α*_1_, …, *α*_*m*_) with *α*_*m*_ = 1 for *m* = 1, …, *M*. The naive observer posterior 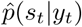 simply updates the flat prior based on only the most recent observation *y*_*t*_. Hence, we have 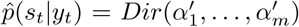 with 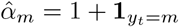.

A detailed account of the readout definitions can be found in S1 Appendix and exemplary time courses of the readouts of the DC model are depicted in Fig 6.

**Fig 6.**
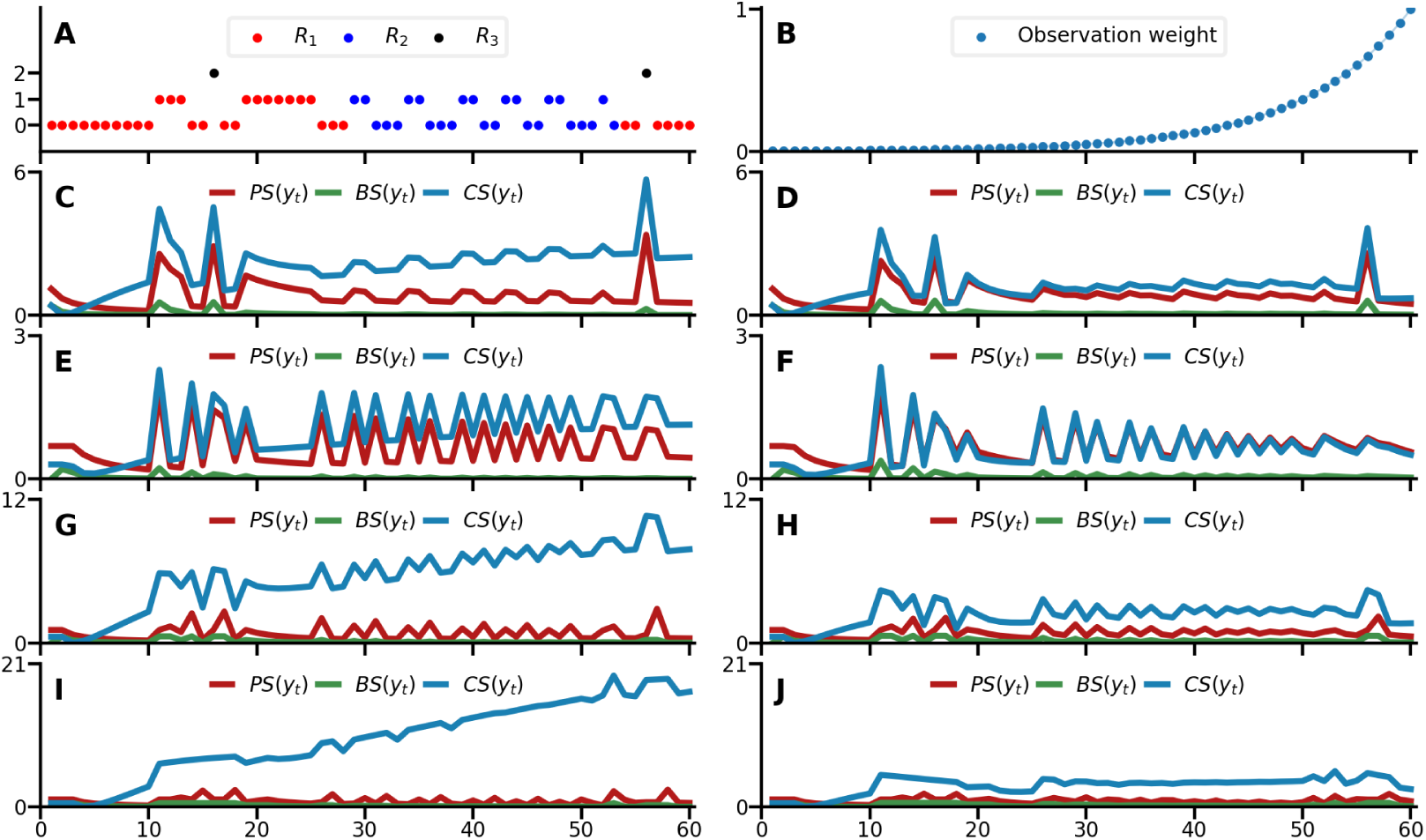
Surprise readouts of the Dirichlet-Categorical model. A) Example sequence with *o*_*t*_ = 0, *o*_*t*_ = 1, and *o*_*t*_ = 2. The forgetting kinetic belonging to *τ* = 0.1, plotted in B, is discounting past observations. C-J) Surprise readouts from the models are displayed either without forgetting on the left-hand side in C (SP), E (AP), G (TP_1_), and I (TP_2_) and with forgetting for *τ* = 0.1 on the right-hand side in D (SP), F (AP), H (TP_1_), and J (TP_2_).

For the HMM, the surprise readouts are obtained by iteratively computing the posterior distribution via the Baum-Welch algorithm using the *hmmlearn* Python package [57]. For timestep *t* this entails fitting the HMM for a stimulus sequence *o*_1_, …, *o*_*t*_ which gives a set of parameter estimates, 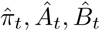 and the filtering posterior 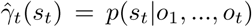. Predictive, Bayesian, and confidence-corrected surprise may then be expressed as follows (see S1 Appendix). An example of the different inference types (SP, AP, and TP) and their surprise readout functions may be found in Fig 7.

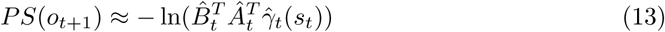

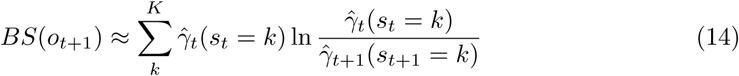

**Fig 7.**
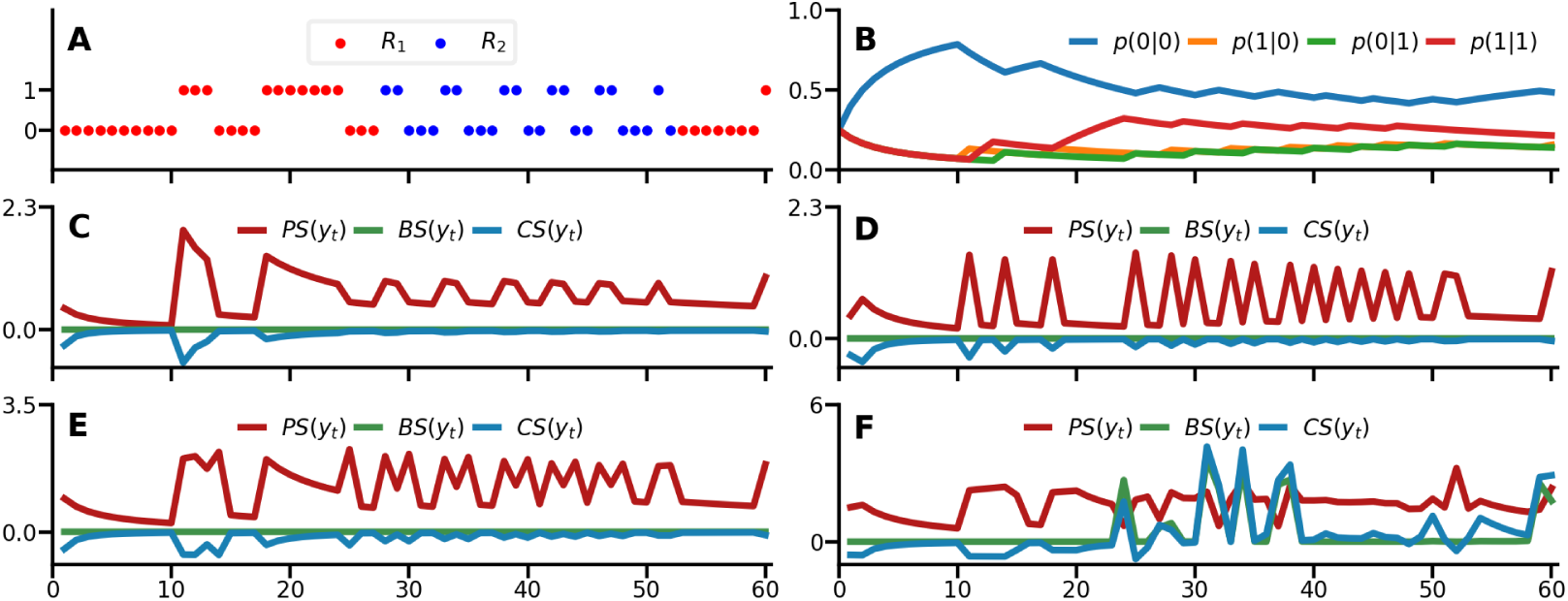
Surprise readouts of the Hidden Markov Model. A) Example sequence with *o*_*t*_ = 0 and *o*_*t*_ = 1. B) Evolution of the estimated emission probabilities of the HMM TP model over time for one of its latent states. C-F) Surprise readouts from the models: SP (C), AP (D), TP_1_ (E), and TP_2_ (F).

Following Faraji et al. [35], confidence-corrected surprise may be expressed as a linear combination of predictive surprise, Bayesian surprise, a model commitment term (negative entropy) *C*(*p*(*s*_*t*_)), and a data-dependent constant scaling the state space *O*(*t*). Here we make use of this alternative expression of CS in order to facilitate the HMM implementation:

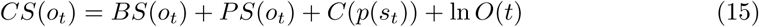

### Model fitting via free-form variational i nference a lgorithm

Each combination of model class (DC and HMM), inference type (SP, AP, TP_1_, TP_3_), and surprise readout function (PS, BS, CS) yields a stimulus sequence-specific regressor. These regressors, as well as those of a constant null-model, were fitted to the single-trial, event-related electrode and source activation data. Using a free-form variational inference algorithm for multiple linear regression [58–60], we obtained the model evidences allowing for Bayesian model selection procedures [61], which accounts for the accuracy-complexity trade-off in a formal and well established manner [62]. In short, the single-subject, single peri-stimulus time bin data *y* ∈ ℝ^*n*×1^ for *n* ∈ *N* trials was modeled in the following form:

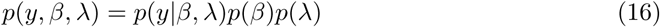

with *β* ∈ ℝ^*p*^ and *λ* > 0 denoting regression weights and observation noise precisions, respectively. The parameter-conditional distribution of *y, p*(*y*|*β, λ*), is specified in terms of a multivariate Gaussian density with expectation parameter *Xβ* and spherical covariance matrix. The design matrix *X* consisted of a constant offset (null-model: *X* ∈ ℝ^*n*×1^) and an additional surprise-model specific regressor in case of the non-null models (*X* ∈ ℝ^*n*×2^). Both a detailed description of the algorithm and the test procedure performed on simulated data used to select the prior parameters for the variational distributions of *β* and *λ* may be found in the supplementary material.

### Bayesian model selection

Before modeling single subject, single peri-stimulus time bin data (*y*) as described above, the single-trial regressors of all non-null models as well as the data underwent z-score normalization to allow for the use of the same model estimation procedure for both sensor and source data. For single subjects, data and regressors corresponding to the five experimental runs were concatenated prior to fitting. To allow for the possibility that the brain estimates statistics computed across multiple timescales of integration [9, 63, 64], the forgetting-parameter *τ* of the DC model was optimized for each subject, model, and peri-stimulus time-bin. To this end, DC model regressors were fitted for a logarithmically spaced vector of 101 *τ*-values on the interval of 0 to 1 and the value of *τ* that resulted in the highest model evidence was chosen. To penalize the DC model for having one of its parameters optimized, the degree to which *τ* optimization on average inflated model evidences was subtracted prior to the BMS procedure.

The furnished model evidences were subsequently used for a random-effects analysis as implemented in SPM12 [61] to determine the models’ relative performance in explaining the EEG data. In order to combat the phenomenon of model-dilution [65], a hierarchical approach to family model comparison was applied (for a graphical overview see S3 Fig). Note that this procedure is performed for each peri-stimulus time bin independently. In a first step, the two model classes DC and HMM were compared against each other and the null-model in a family-wise BMS. A threshold of exceedance probabilities *ϕ*>0.99 in favour of either the DC or HMM was applied, so that only whenever there was very strong evidence in favour of one of the model classes the following analyses were applied. For timepoints with exceedence probabilities above this threshold, a family-wise comparison of TP_1_ and TP_2_ was performed in order to determine which order of transition probabilities would be used for the second level. Subsequently, either the TP_1_ or TP_2_ models were compared to the SP and AP models. Wherever *ϕ*>0.95 for one of the inference type families, the third analysis level was called upon. On this final level, surprise read-out functions were compared for the winning model class and corresponding inference type with a threshold of *ϕ*>0.9. As such, this step-wise procedure allows spatio-temporal inference of the read-out functions for which there is strong evidence of the belonging model class and inference type. The same procedure was used for the EEG sensor and source data.

To inspect the values of the forgetting-parameter *τ* that best fit the data, subject specific free energy values were averaged across the timebins with significant surprise readout effects for the corresponding dipoles. These were summed across subjects to yield the group log model evidence for each tested value of *τ*, which were subsequently compared against each other.

## Results

### Behavioural results and event-related potentials

Participants showed consistent performance in counting the amount of catch trials during each experimental run, indicating their ability to maintain their attention on the stimuli (robust linear regression of presented with reported targets: slope=0.96, p<0.001, R^2^=0.93). Upon questioning during the debriefing, no subjects reported explicit awareness of switching regimes during the experiment.

An initial analysis was performed to confirm our paradigm elicited the typical somatosensory responses. Fig 8B shows the average SEP waveforms for contralateral (C4, C6, CP4, CP6) somatosensory electrodes with the expected evoked potentials, i.e. N20, P50, N140 and P300 resulting from stimulation of the left wrist. The corresponding topographic maps (Fig 8C) confirm the right lateralized voltage distribution of the somatosensory EEG components on the scalp. The EEG responses to stimulus mismatch were identified by subtracting the deviant from the standard trials (deviants-standards), thereby obtaining a difference wave for each electrode (see Fig 8D). The scalp topography of the peak differences between standards and deviants within predefined windows of interest indicates mismatch responses over somatosensory electrodes (Fig 8E).

**Fig 8.**
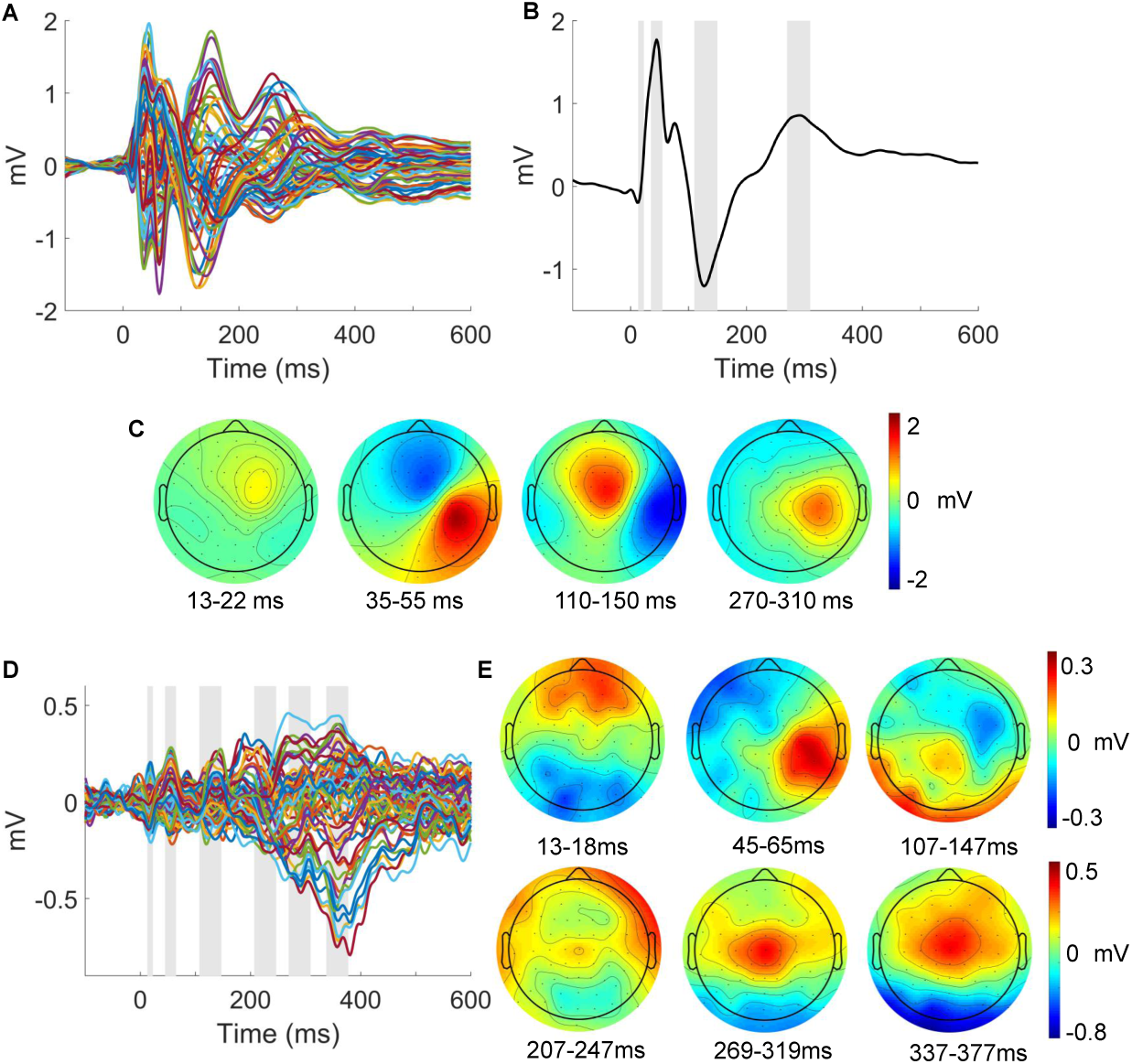
Event-related potentials. A) Grand average SEP of all 64 electrodes. B) Average SEP across electrodes C4, C6, CP4, CP6 (contralateral to stimulation). Grey bars indicate time windows around the standard somatosensory ERP components (13-23ms; 35-55ms; 110-150ms; 270-310ms). C) ERP scalp topographies corresponding to the time windows in B. D) Grand average ERP of the mismatch response obtained by subtraction of standard from deviant trials of 64 electrodes. Grey bars indicate windows around peaks which were identified within pre-specified time windows of interest around somatosensory ERP or expected mismatch response components (13-18ms; 45-65ms; 107-147ms; 207-247ms 269-319ms; 337-377ms). ERP scalp topographies corresponding to the time windows in D)

To test for statistical differences in the EEG signatures of mismatch processing we contrasted standard and deviant trials with the general linear model. Three main clusters reached significance after performing family-wise error correction for multiple comparisons. The topographies of resulting F-values are depicted in Fig 9. The earliest significant difference between standard and deviant trials can be observed around 60ms post-stimulus (peak at 57ms, closest electrode CP4, p_FWE_=0.002, F=27.21, Z=5.07), followed by a stronger effect of the hypothesized N140 component around 120ms which will be referred to as the N140 mismatch response (N140 MMR, peak at 119ms, closest electrode: FC4, p_FWE_=0.001, F=29.56, Z=5.29). A third time window of a very strong and elongated difference effect starting around 250ms to 400ms post-stimulus which corresponds to the hypothesized P300 MMR (peak at 361ms, closest electrode: Cz, p_FWE_<0.001, F=72.25).

**Fig 9.**
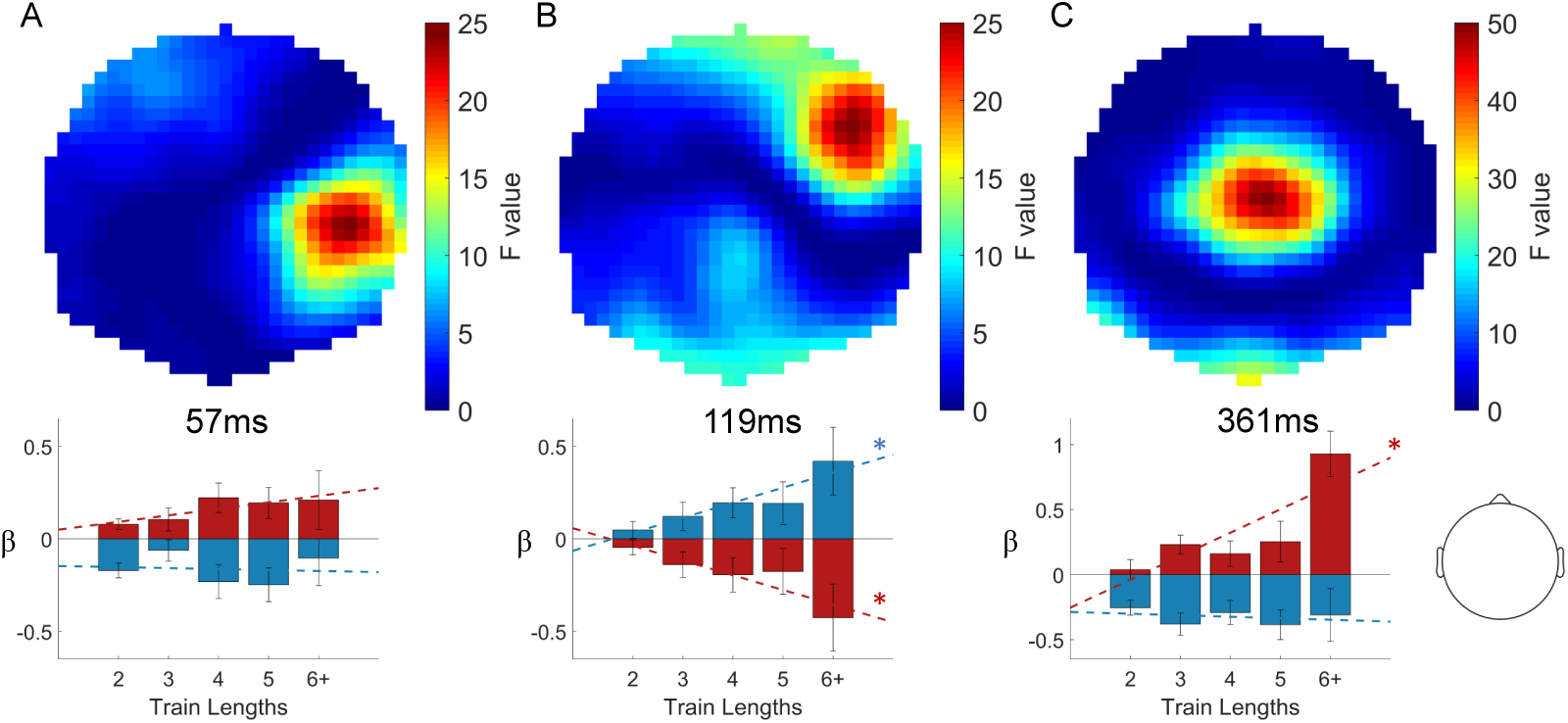
Statistical parametric maps of mismatch responses. Top row: Topographical F maps resulting from contrasting standard and deviant conditions averaged across the times of significant clusters: 57ms (A), 119ms (B) and 361ms (C). Bottom row: Corresponding beta parameter estimates of the significant peaks. Asterisks indicate significant linear fits (p<0.05). Head depiction on the bottom right shows the orientation of the topographic maps.

The inspection of the *β*-parameter estimates at the reported GLM cluster peaks (illustrated in Fig 9) indicates that stimulus train length, i.e. the number of standard stimuli that precede a deviant stimulus, has differentiable effects on the size of EEG responses to standard and deviant stimuli. Both the N140 and P300 MMR effects are found to be parametrically modulated by train length as indicated by a significant linear relationship between *β*-estimates and train length. Specifically, the N140 MMR effect is reciprocally modulated by stimulus type, such that responses to standards are more positive for higher train lengths (F-statistic vs. constant model: 5.45, p=0.021) while deviant responses become more negative (F-statistic vs. constant model: 5.07, p=0.026). The parametric effect on the P300 MMR is entirely driven by the effect on deviant stimuli (F-statistic vs. constant model: 20.7, p<0.001), with no effect of train length on the response to standard stimuli (p>0.05). For the early 60ms cluster no effect was found on either standard or deviant stimuli.

### Source Reconstruction

The distributed source reconstruction resulted in significant clusters at the locations of primary and secondary somatosensory cortex (Fig 10A, with details specified in the corresponding table). The resulting anatomical locations were subsequently used as priors to fit four equivalent current dipoles (Fig 10B, with details specified in corresponding table). Two dipoles were used to model S1 activity at time points around the N20 and the P50 components while an additional symmetric pair captured bilateral S2 activity around the N140 component.

**Fig 10.**
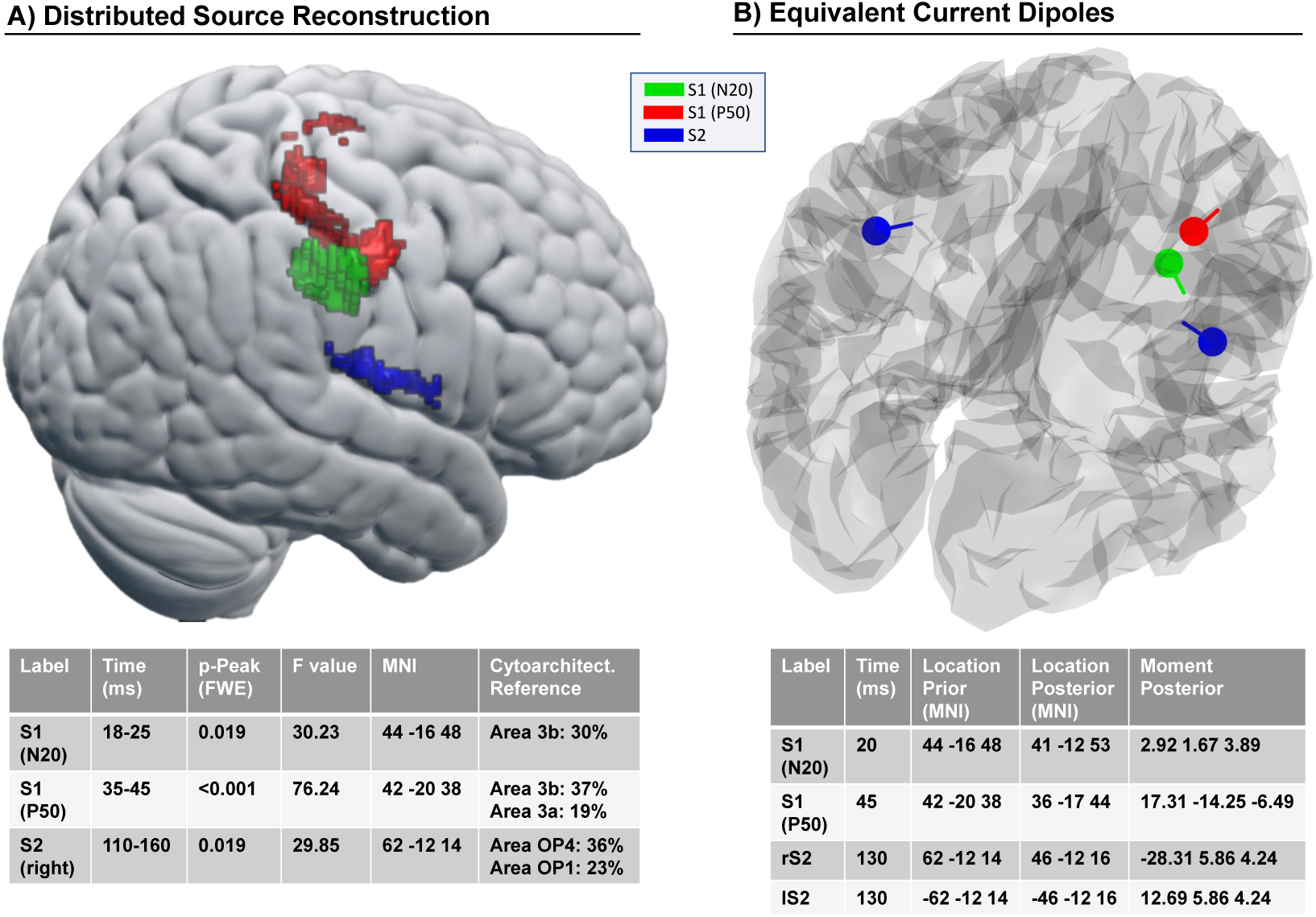
EEG source model. A) Statistical results of distributed source reconstruction. Red: 18-25ms, Green: 35-45ms, Blue: 110-160ms. Below: Table with corresponding detailed data of the clusters. B) Location and orientation of fitted equivalent current dipoles. Red: S1 (N20), Green: S1 (P50), Blue: bilateral S2. Below: Table with their corresponding values.

To establish the plausibility of the somatosensory dipole model the EEG data was projected onto the four ECD’s and the grand average source ERP was computed across subjects for standard and deviant trials. The resulting waveforms, shown in Fig 11, show a neurobiologically plausible spatiotemporal evolution: the two S1 dipoles reflect the early activity of the respective N20 and P50 components while the S2 dipoles become subsequently active and show strongest activity in right (i.e. contralateral) S2. The average response to standards and deviants within time windows around the significant MMRs in sensor space (around 57ms and 119ms; see Fig 9) were compared with simple paired t-tests. The S1_P50_ dipole shows a significant difference at both time windows and can be suspected to be the origin of the effect at 57ms as well as contribute to the 119ms MMR (both p<0.05; bonferroni corrected) while the right S2 dipole is mainly driving the strong 119ms effect (p<0.01; bonferroni corrected).

**Fig 11.**
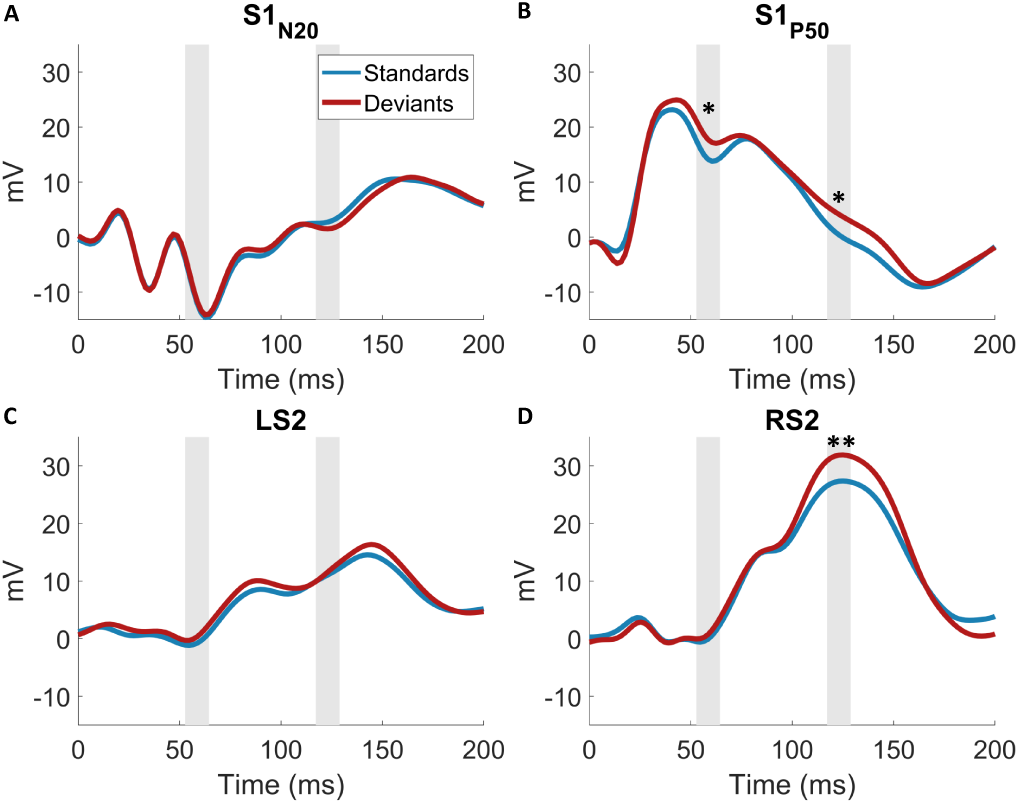
Grand average waveforms of EEG dipole projections. Standards and deviants were contrasted within time windows of interest informed by the GLM in the results section. *p<0.05; **p<0.01; Bonferroni corrected

### Single trial modeling

We previously established the presence of mismatch responses in sensor space and confirmed their origin in the somatosensory system by modeling the early EEG components in source space. Subsequently, we investigated the temporal and spatial surprise signatures with trial-by-trial modeling of electrode and source data.

### Modeling in sensor space

For large time windows at almost all electrodes there is strong evidence in favor of the DC model class (*ϕ*>0.99), while the HMM model class does not reach significance anywhere, therefore excluding HMM models from further analyses. To verify that this result was not merely due to an insufficient penalization of the DC models, the analysis was repeated with *τ* =0. Thus, under this setting, all instances of the DC model had perfect, global integration similar to the HMM models. Likewise, no significant results were found for the HMM model class. For the DC model, TP_1_ is found to outperform TP_2_, thus excluding TP_2_ for the second and third level analyses. In the following step, TP_1_ clearly performed better than SP and AP at almost all electrodes and time points (see Fig 12A-C). Thus, the results reported in the following section focus on the comparison of the readout functions of the DC TP_1_ model.

**Fig 12.**
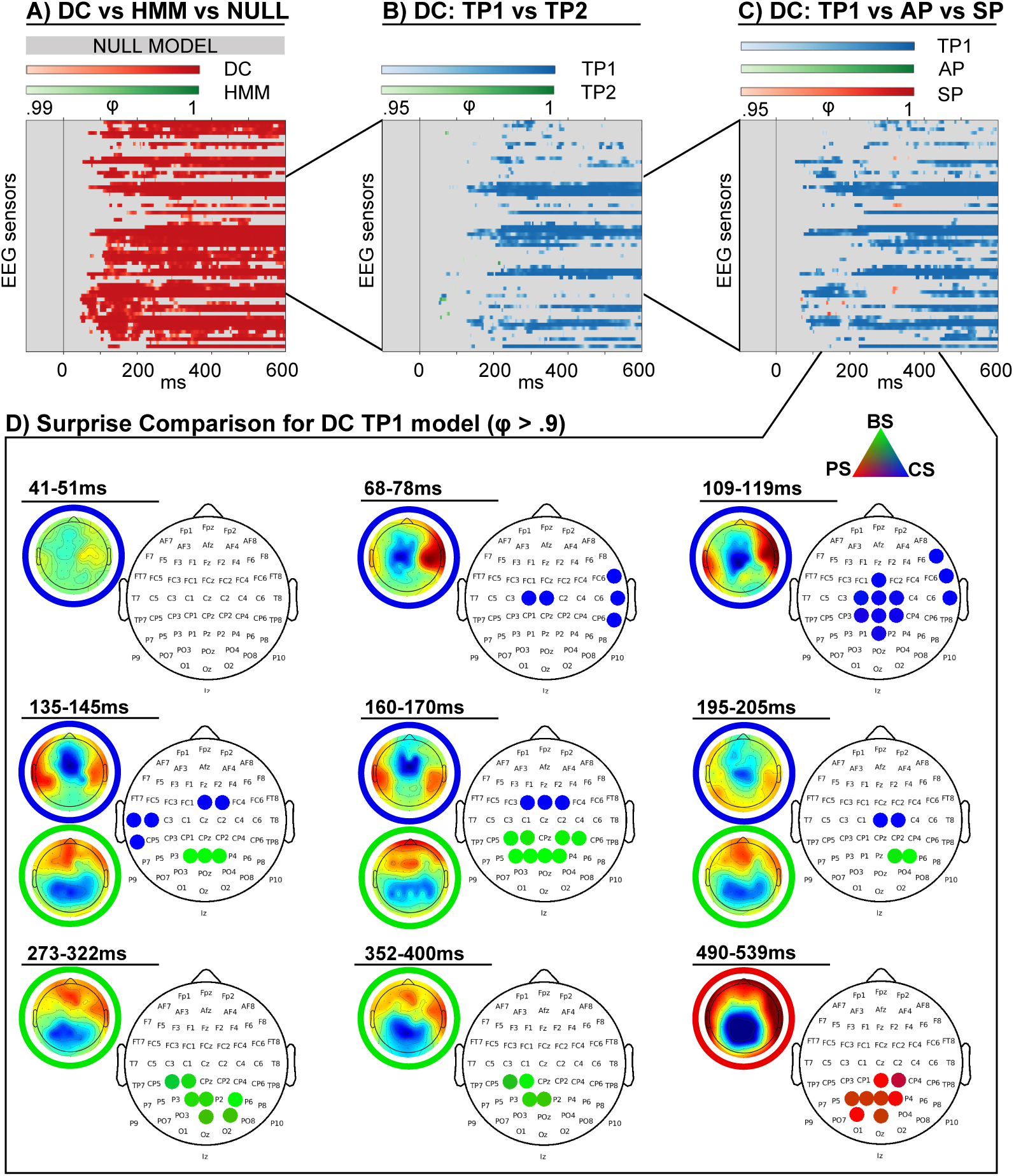
Modeling results. Exceedence probabilities (*φ*) resulting from the RFX family model comparison. A) Dirichlet-Categorical (DC) model, Hidden Markov Model (HMM) and Null model family comparison, thresholded at *φ*>0.99:. B) Family comparison within the winning DC family, thresholded at *φ*>0.95: first and second order transition probability models (TP1, TP2). C) Family comparison within the winning DC family, thresholded at *φ* > 0.95: first order transition probability (TP1), alternation probability (AP) and stimulus probability (SP) models. D) Family comparison of surprise models within the winning DC TP1 family, thresholded at *φ*>0.9: Large discrete topographies show the significant electrode clusters of Predictive surprise (PS) in red, Bayesian surprise (BS) in green and confidence-corrected surprise (CS) in blue. Small continuous topographies display the converged variational expectation parameter *m*_*β*_.

Fig 12D shows the result of the random-effects Bayesian model selection analysis. The scalp topographies depict the winning readout functions of the Dirichlet-Categorical TP model at different time windows. The converged variational expectation parameter *m*_*β*_ resulting from the model fitting procedure (see S2 Appendix) are displayed for the winning models to facilitate interpretation of the topography. Given the difference in temporal dynamics of faster, early (<200 ms) and slower, late (200-600ms) EEG components, different thresholds were applied. Early significant clusters were identified by averaging exceedance probabilities over 10ms windows and using a minimum cluster size of two electrodes. After 200ms, clusters were identified by averaging over 50ms time windows with a minimum cluster size of four. From around 70ms on, early surprise computations can be observed with confidence corrected surprise (CS) best explaining the EEG data on contralateral and subsequently ipsilateral electrodes up to around 200ms. A significant cluster of Bayesian surprise (BS) is prominent at centro-posterior electrodes between 130-200ms, with similar electrodes later again representing Bayesian surprise around 300 and 375ms. These clusters are temporally in accordance with the N140 and P300 MMR effects. The latest cluster at around 500ms post-stimulus is entirely driven by predictive surprise.

### Modeling in source space

The topographic distribution of the effects of confidence-corrected surprise seem to indicate an early contribution of secondary somatosensory cortex from around 70ms on that starts contra- and extends ipsilaterally while the BS cluster emerges around the time of the N140 MMR. In order to further investigate this interpretation and examine the spatial origin of the 140-190ms Bayesian surprise cluster, we fit our models to the single trial dipole data and used the same hierarchical Bayesian model selection approach as for the sensor-space analysis described in the methods section. Results for the source activity were highly similar, with significant results exclusively for the DC TP_1_ model. Consequently, the surprise readout functions of the TP_1_ model were subjected to BMS. The results depicted in Fig 13 confirm the early onset of CS in secondary somatosensory cortex and allocate the later onset BS cluster to primary somatosensory cortex. Additionally, a significant PS effect with an onset of around 170ms in the right S2 dipole is observed, which did not reach significance in the sensor-level analysis.

**Fig 13.**
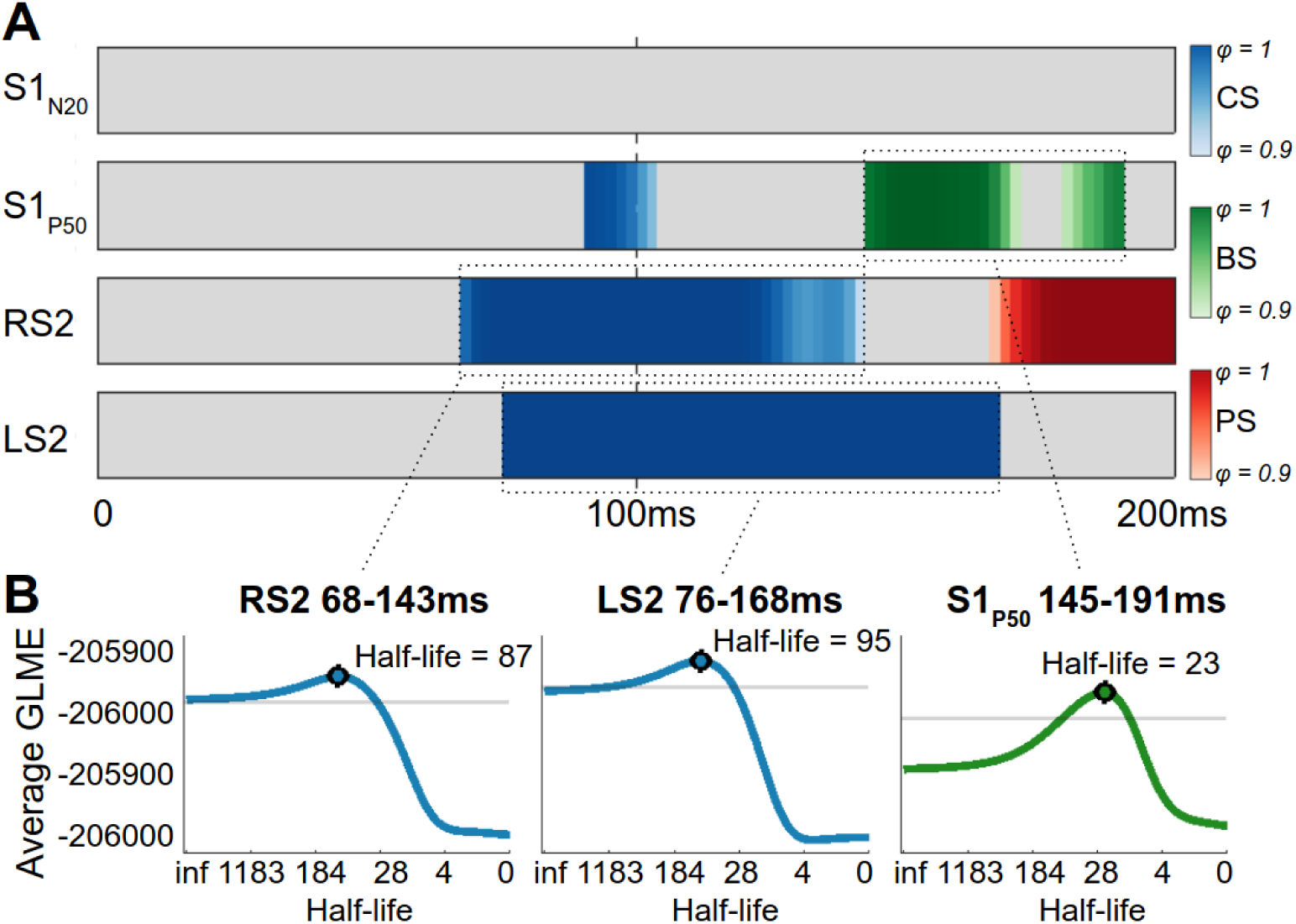
Modeling results in source space with best fitting forgetting-parameter values. Red: Predictive surprise (PS), Green: Bayesian surprise (BS), Blue: Confidence-corrected surprise (CS) A) Colors depict significant time points for the surprise readout functions of the Dirichlet-Categorical TP_1_ model within the dipoles S1_N20_, S1_P50_, right S2 (RS2) and left S2 (LS2). B) The group log model evidence (GLME) values corresponding to the stimulus half-lives for forgetting-parameter *τ*, after averaging the significant timebins for the right S2 (RS2), left S2 (LS2), and S1_P50_ dipoles. The grey lines indicate a difference of 20 GLME from the peak, indicating very strong evidence in favour of the peak half-life value compared to values below this threshold.

### Leaky integration

We inspected the *τ* -parameter values that resulted in the highest group log model evidence for the reported significant dipole effects (Fig 13). All three considered clusters indicate a local timescale of integration, with the best-fitting parameter values resulting in a stimulus half-life of ∼95 and ∼87 for the confidence-corrected surprise effects at 68-143ms and 76-168ms respectively, and a half-life of ∼23 stimuli for the Bayesian surprise effect at 145-191ms. Using the single-subject peaks, *τ* was found to significantly differ from 0 (i.e., no forgetting) for the BS effect in S1 (p<0.001) and CS in RS2 (p<0.05), but not in LS2 (p=0.06). Paired t-tests revealed no significant difference in *τ* underlying the three effects (p>0.05).

## Discussion

In this study, we used a roving paradigm to identify EEG mismatch responses independent of stimulus identity. Two early effects were source localized to the somatosensory system and indicated change detection and perceptual learning mechanisms, while a late P300 effect was characteristic of attention-allocating processes. Using computational modelling, EEG signals were best described using a non-hierarchical Bayesian learner performing transition probability inference. Furthermore, we dissociated an early representation of confidence-corrected surprise localized to bilateral S2 with subsequent Bayesian surprise encoding in S1. These computations were shown to use a local, rather than global, timescale of integration.

We report a significant somatosensory mismatch response around three distinct post-stimulus time-points: 57ms, 119ms, and 361ms. These will be referred to as sMMR’s as opposed to MMN since the effects at 57ms and 361ms are not negativities and our experimental protocol included an explicit attentional focus on the stimulation. The MMN was originally defined to be a pre-attentive effect and while attention to the stimulus does not seem to influence the MMN in the visual domain [66], we don’t address a potential independence of attention here. Nevertheless, the reported sMMR effects integrate well with previous findings on the somatosensory MMN (sMMN). Our 119ms effect is in line with the timing of the most commonly reported sMMN as a modulation of the N140 component between 100-250ms [17, 27, 24, 21–23, 30, 28, 31, 29, 25, 26]. However, some studies additionally describe a modulation of multiple somatosensory components [17–19, 24], similar to our 3 distinct sMMR effects. The electrode positions reported in sMMN studies show a large variability of frontocentral and parietal electrodes. These discrepancies might be driven by the differences in stimulation sites (different fingers and hand) and deviant definitions (vibrotactile frequencies, stimulation site, stimulation duration). Here, we present significant effects around C4 and FC4 electrodes for the 57 and 119ms time-points, respectively, indicating EEG generators within the contralateral somatosensory system. This implication is in line with intracranial EEG recordings of the somatosensory cortex during oddball tasks [24, 30]. In accordance with previous MEG studies using source localization [21, 22], our source space analysis suggests the early MMR effects to originate from contralateral primary and secondary somatosensory cortex (cS1 and cS2, respectively), with the earliest MMR (at 57ms) localized to cS1 followed by a combined response of S1 and S2. While evidence exists for a role of S2 in the early phase of mismatch processing [26], the evolution from an initial MMR generated by S1 to an additional involvement of S2 in the mid-latency MMR, as indicated by our findings, is consistent with the sequential activation of the somatosensory hierarchy in general tactile stimulus processing [67–69]. Finally, the third sMMR effect at 361ms is in accordance with a large body of evidence showing a modulation of the P300 component by mismatch processing [70–72]. The P300 in response to oddball tasks likely reflects a modality unspecific effect, dependent on task-related resource allocation [73–77] and contingent on attentional engagement [29]. In this regard, our observed late sMMR effect is related to the subjects’ attentional focus on the stimulus sequence and, more specifically, might reflect an updating process of the attention-allocating mechanism as suggested by Kopp et al. [41].

In addition to three spatiotemporally distinct sMMR effects, we further show their differential modulation by the length of standard stimulus trains preceding the deviant stimulus. This finding supports the interpretation that distinct mechanisms underlie the generation of the different sMMR’s. The earliest effect around 57ms is not affected by train length, likely reflecting a basic change detection mechanism that signals a deviation from previous input regardless of statistical regularities. The mid-latency MMR around 119ms, on the other hand, shows a significant linear dependence on stimulus train length for both deviant and standard stimuli. Longer train lengths result in parametrically stronger negative responses to deviant stimuli while responses to standard stimuli are increasingly reduced. This effect is in accordance with repetition suppression effects reported for the MMN [78, 79] which have been shown to be dependent on sequence statistics and are interpreted to reflect perceptual learning [80, 81]. While it has been indicated that the number of preceding standards can also enhance the sMMN [26], no previous studies show comparable effects to our parametric modulation of the mid-latency sMMR. The reciprocal effect of repetition for standard and deviant stimuli shown here indicate early perceptual learning mechanics in the somatosensory system, likely originating from S2 in interaction with S1. In contrast, later mismatch processing reflected by the sMMR at 361ms only shows a linear dependence of deviant stimuli on train length, while the response to standards remains constant. This is in line with the interpretation that perceptual learning in the P300 reflects a recruitment of attention in response to environmental changes, possibly accompanied by updates to this attentional-control system [41].

In addition to average-based ERP analyses, single-trial brain potentials in response to sequential input can provide a unique window into the mechanisms underlying probabilistic inference in the brain. Here, we investigated the learning of statistical regularities using different Bayesian learner models with single-trial surprise regressors. Partitioning the model space allowed us to infer on distinguishing features between the model families using Bayesian model selection (BMS). The first comparison concerned the form of hidden state representation: In order for a learner to adequately adapt one’s beliefs in the face of changes to environmental statistics, more recent observations may be favored over past ones without modeling hidden state dynamics (Dirichlet-Categorical model; DC), or different sets of statistics may be estimated for a discretized latent state (Hidden Markov Model; HMM). Our comparison of these two learning approaches provides strong evidence for the DC model class over the HMM for the large majority of electrodes and post-stimulus time. The superiority of the DC model was found to be irrespective of the inclusion of leaky integration to the DC model, indicating the advantage of a non-hierarchical model in explaining the EEG data. Even though the data generation process included discrete hidden states in the form of fast and slow switching regimes, participants were neither aware of their existence nor was their dissociation required to perform the behavioural task. As such, the early EEG signals studied here are likely to reflect a form of non-conscious, implicit learning of environmental statistics [82–84]. However, it is possible that the brain implements different learning algorithms in different environments, resorting to more complex ones only when the situation demands it. Indeed, humans seem to assume different generative models in different contexts, possibly depending on task instructions [85]. This may in part explain why evidence has been provided for the use of both hierarchical [86, 87] and non-hierarchical models [88, 89]. By omitting instructions to learn the task-irrelevant statistics, our study potentially avoids the issue of invoking a certain generative model. We might therefore report on a ‘default’ model of the brain used to non-consciously infer environmental statistics.

In order to investigate which statistics are estimated by the brain during the learning of categorical sequential inputs, we compared three models within the DC model family that use different sequence properties to perform inference on future observations: stimulus probability (SP), alternation probability (AP), and transition probability (TP) inference. The TP model subsumes SP and AP models and is thus more general by maintaining a larger hypothesis space. Our results show that the TP model family clearly outperformed the SP and AP families, thereby suggesting that the brain captures sequence dependencies by tracking transitions between types of observations for future inference. We thereby provide further evidence for an implementation of a minimal transition probability model in the brain as recently concluded from the analysis of several perceptual learning studies [90], extending it to include somesthesis. Additionally, we expand upon previous studies by comparing a first order TP model (TP_1_), capturing transitions between stimuli conditional only on the previous observation, with a second order TP model (TP_2_), which tracks transitions conditional on the past two observations. Our results suggest that the additional complexity of the second order dependencies contained in our stimulus sequence were not captured by the brain, although it may resort to alternative, more compressed representations [91].

The BMS analyses of the partitioned model space suggests that the brain’s processing of the stimulus sequences is best described by a Bayesian learner with a static hidden state (akin to the DC model) which estimates first-order transition probabilities (TP1). Within the DC-TP1 model family, we compared the surprise quantifications themselves as the readout functions for the estimated statistics of the Bayesian learner: predictive surprise (PS), Bayesian surprise (BS), and confidence-corrected surprise (CS). The results show that the first surprise effect is represented by CS from around 70ms over contralateral somatosensory electrodes which rapidly extends bilaterally around 80ms, dissipating around 170ms. BS is found as a second significant centro-posterior electrode cluster of surprise between 140-190ms. As proposed by Faraji et al. [35], CS is a fast-computable measure of surprise in the sense that it may be computed before model updating occurs. In contrast, as BS describes the degree of belief updating, which requires the posterior belief distribution, it is expected to be represented only during the update step or later. As such, the temporal evolution of the observed CS and BS effects is in accordance with the computational implications of these surprise measures. Specifically, our study provides evidence for the hypothesis that the representation of CS, as a measure of puzzlement surprise, precedes model updating and may serve to control update rates. While PS is also a fast-computable puzzlement surprise measure and (similarly to CS) is scaled by the subjective probability of an observation, CS additionally depends on the confidence of the learner, read out as the (negative) entropy of the model. Evidence for a sensitivity to confidence of prior knowledge in humans has been reported in a variety of tasks and modalities [92–94]. It is important to note that one may also be confident about novel sensory evidence (e.g. due to low noise) which may result in larger model updates [95]. This aspect of confidence, however, lies outside the scope of the current work and it will be left to future research to investigate its role in somatosensory learning.

Our source reconstruction analyses attributed the early CS effects to the bilateral S2 dipoles, which is in accordance with the timing of S2 activation reported in the literature [67–69]. This finding suggests that the secondary somatosensory cortex may be involved in representing confidence about the internal model. The BS effect around 140ms was localized to S1 in source space, with its timing matching prior work using Bayesian modeling of surprise signals in the somatosensory system [13]. Additionally, our findings are in accordance with previous accounts of perceptual learning in the somatosensory system [96]. It should be noted that even though the BMS performed in source space shows an additional PS representation at the right-S2 dipole from 170ms onward, the effect is mainly driven by the activity of two electrodes within a short time window and does not survive thresholding in the sensor level analysis. Thus, it is likely that our restricted dipole analysis is not appropriately source localizing these effects as they might originate from diffuse sources not included in our model. Such effects in source space at the edge of our analysis window must therefore be taken with caution and we refrain from interpreting them here. In sum, these results suggest that the secondary somatosensory cortex may represent confidence about the internal model and compute early surprise responses, potentially controlling the rate of model updating. Signatures of such belief updating, on the other hand, were found in S1 around the time of the N140 somatosensory response. These effects together indicate a possible interaction between S1 and S2 that is responsible for both a signaling of the inadequacy of the current beliefs and their subsequent updating.

Our implementation of the Dirichlet-Categorical model incorporates a memory-decay parameter *τ* that exponentially discounts observations in the past. The *τ*-values for the winning models of our BMS analyses that best fit the data within the significant spatiotemporal clusters indicate relatively short integration windows for both CS and BS with stimulus half-lives of approximately 85 and 25 observations, respectively. This suggests that, within our experimental setup, the brain uses local sequence information to infer upcoming observations rather than global integration, for which all previous observations are considered. For a sub-optimal inference model with a static hidden state representation, the incorporation of leaky observation integration on a more local scale can serve as an approximation to optimal inference resorting to dynamic latent state representation and can thereby capture a belief about a changing environment [90].

Given a very large timescale, BS converges to zero as the divergence between prior and posterior distributions decreases over time, imposing an upper bound on the timescale. Meanwhile, for PS and CS it tends to lead to more accurate estimates of p(y_*t*_|s_*t*_) as more observations are considered. However, given the regime switches in our data generation process, a trade-off exists where a timescale that is too large prevents flexible adaptation following such a switch. In the current context, the timescales are local enough where the estimated statistics are able to be adapted in response to regime switches (with a switch occurring every 100 stimuli on average). Especially CS shows a large range of *τ*-values producing similarly high model evidence due to the high correlation between regressors. As no significant differences in optimal *τ*-values between clusters were found, it might be the case that the same timescale is used for the computation of both CS and BS. Although the uncertainty regarding the exact half-lives is in line with the large variability found in the literature, local over global integration is consistently reported [13, 39, 90, 9, 48, 91]. Given a fixed inter-stimulus interval of 750ms, a horizon of 85 and 25 observations may be equated to a half-life timescale of approximately 63 to 17 seconds, with regime switches expected to occur every 75 seconds.

Some considerations of the current study deserve mention. First, the behavioural task required participants to make a decision about the identity of the stimulus so as to identify target (catch) trials. Thus, one may wonder to what extent the results contain conscious decision making signals, rather than implicit, non-conscious learning activity. However, decision making-related signals are described to occur relatively late in the trial [97, 98] and we assume to largely avoid them here by focusing on early signals prior to 200ms. Secondly, a large model space of both hierarchical as well as non-hierarchical Bayesian learners exists. As such, it is possible that the brain resorts to some hierarchical representation different from the ones tested here. We chose to use an HMM as it closely resembled the underlying data structure, offers the optimal solution for a discrete state environment, and contributes to the field as it has seen only limited application for probabilistic perceptual learning.

In conclusion, we report evidence that early somatosensory processing seems to reflect Bayesian perceptual learning. The system appears to capture a changing environment using a static latent state model that integrates evidence on a local, rather than global, timescale and estimates transition probabilities of observations using first order dependencies. In turn, we show that the estimated statistics are used to compute a variety of surprise signatures in response to new observations, including both puzzlement surprise scaled by confidence (CS) in secondary somatosensory cortex and enlightenment surprise (i.e. model updating; BS) in primary somatosensory cortex.

## Author contributions

SG & MG: Conceptualization, Data curation, Investigation, Project administration, Formal Analysis, Validation, Methodology, Software, Visualization, Writing – original draft, Writing – review & editing

RTL: Conceptualization, Methodology, Software, Visualization, Writing – review & editing

DO: Conceptualization, Methodology, Software, Supervision, Writing – review & editing

FB: Conceptualization, Project administration, Supervision, Funding, Resources, Writing – review & editing

## Supporting information

### S1 Appendix. Bayesian learner models

In this supplementary text we provide the derivations for the presented equations of the compared Bayesian learner models.

### Dirichlet-Categorical model

Given a sequence of observations *y*_1_, …, *y*_*t*_ the Dirichlet-Categorical model combines the likelihood with the prior to refine the posterior estimates over the latent variable space (equation 5):

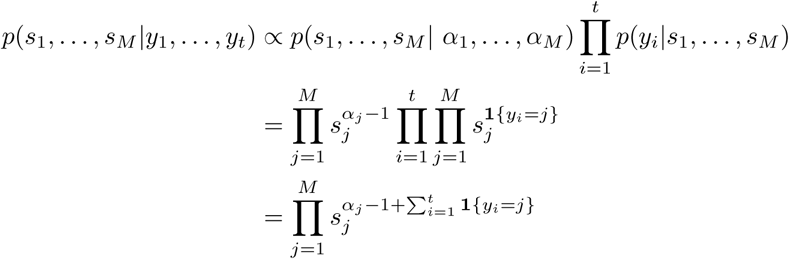

The posterior predictive distribution in equation 7 can be obtained by integrating over the space of latent states:

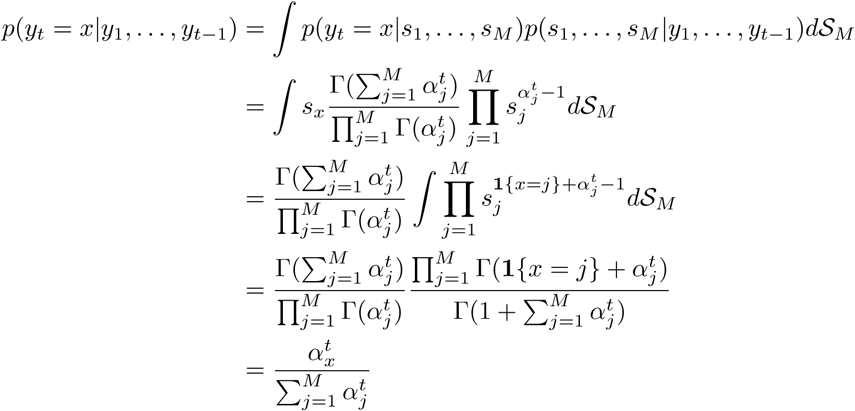

The surprise readout functions for the Categorical-Dirichlet model introduced in equations 10 to 12 are:

#### Predictive Surprise

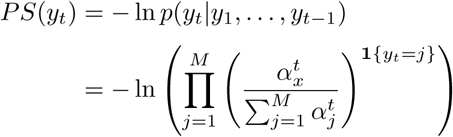

#### Bayesian Surprise

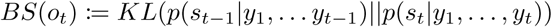

The general KL divergence for two Dirichlet distributions *P* and *Q* parametrized by 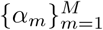 and 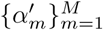 is given by

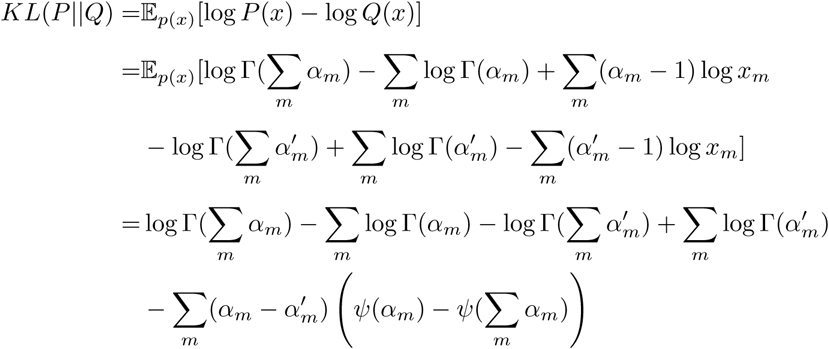

where *ψ*(.) denotes the digamma function.

#### Confidence-Corrected Surprise

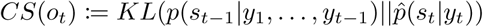

The flat prior can be written as *Dir*(*α*_1_, …, *α*_*m*_) where *α*_*m*_ = 1 for all *m* = 1, …, *M*. The naive observer posterior simply updates the flat prior based on only the most recent observation *y*_*t*_. Hence, we have that 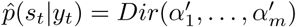 with 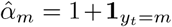. Hence at a given point in time *t*, we have:

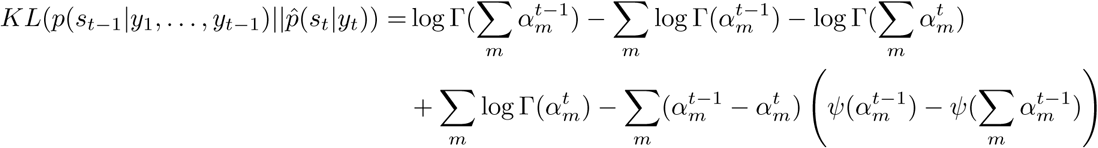

### Hidden Markov Model

For the use of parameter inference via the expectation-maximisation algorithm and in order to derive the factorisiation of the joint likelihood *p*(*o*_1:*t*_, *s*_1:*t*_), we will make sure of the following derivations:

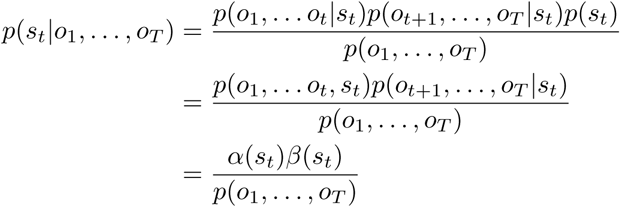

where for the final line we have redefined the backward and forward probabilities as

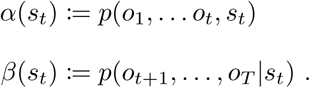

In the following, we derive the forward and backward equations which may be used in conjunction with a Dynamic Programming paradigm such as the Baum-Welch algorithm in order to perform the Expectation-Maximisation inference procedure.

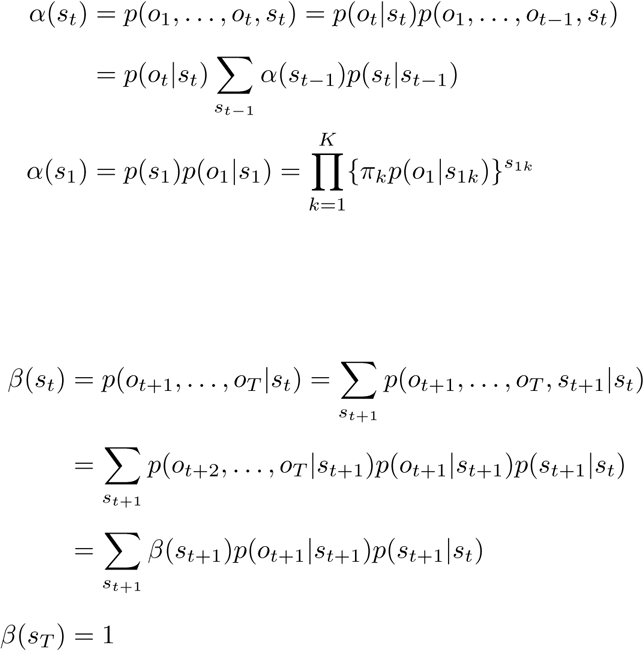

Both expressions for *α* and *β* involve a backward and forward recursion. Given a sequence of observations, these can easily be computed in a sequential fashion. The final quantity of interest for the EM algorithm are the smoothed transition probabilities:

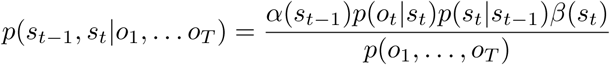

In order to infer the parameters we now alternate between an expectation and a maximisation step:

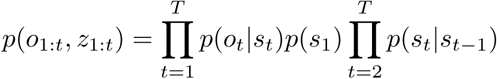

1. **Expectation:**

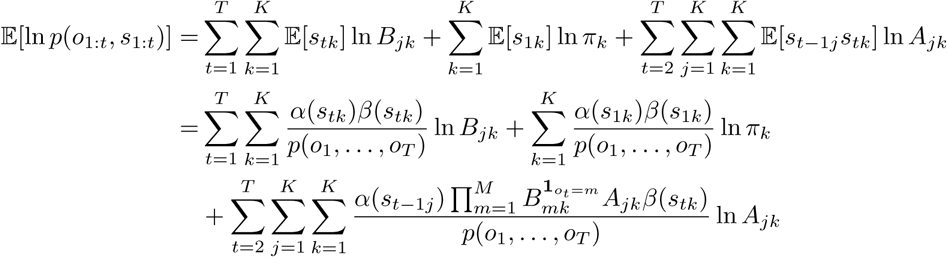
2. **Maximization:** The Lagrangian with the necessary constraints is determined (i.e. row stochasticity and proper distributions) and the derivatives with respect to the set of parameters (*θ* = {*A*_*jk*_, *B*_*mk*_, *π*_*k*_}) is computed.

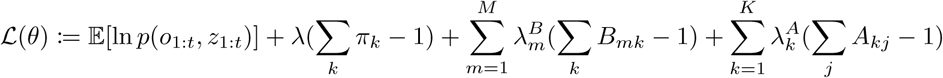

The filtering equation can then be written as

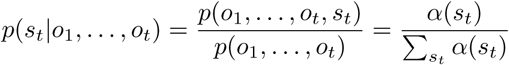

Finally, the evaluation is then easily obtained by marginalising over the hidden state:

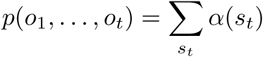

For timestep *t* the HMM was fit for a stimulus sequence *o*_1_, …, *o*_*t*_ which gives a set of parameter estimates, 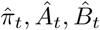 and the filtering posterior 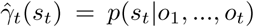. The predictive surprise as formulated in equation 13 is derived in the following way:

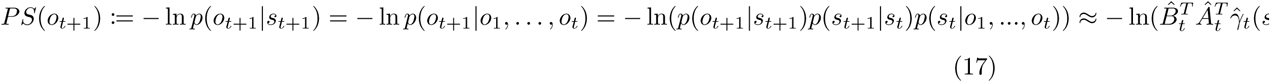

The Bayesian surprise from equation 14, on the other hand, derives for the HMM as follows:

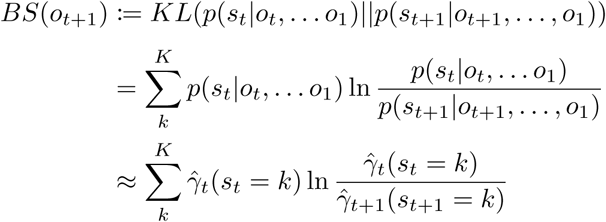

Finally, confidence corrected surprise from equation 15 may be expressed as a linear combination of predictive surprise, Bayesian surprise, a model commitment term (negative entropy) *C*(*p*(*s*_*t*_)), and a data-dependent constant scaling the state space *O*(*t*).

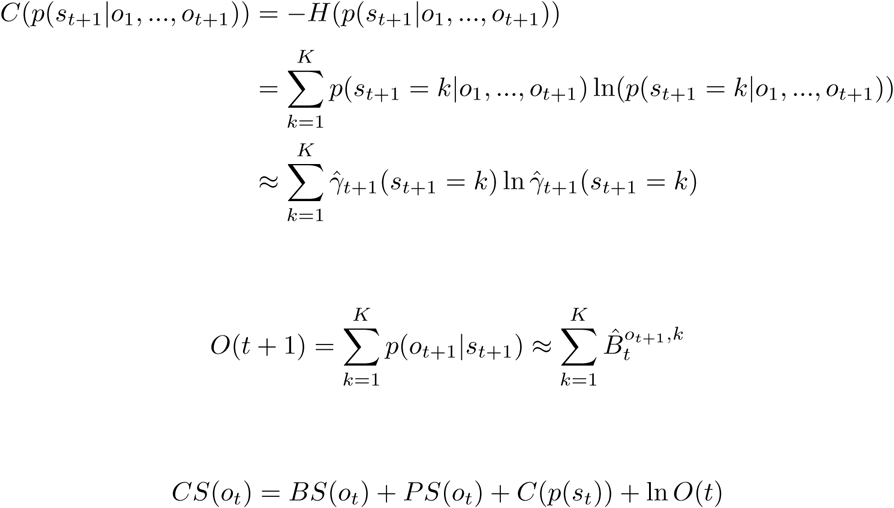

All inference types shared the same state space *s* ∈ *S* = {0, 1}. Due to the transformation of the observation sequence the observation space differed between models:

Stimulus probability model: *y*_*t*_ = *o*_*t*_ for *t* = 1, …, *T* with *𝒪*_*SP*_ = {0, 1}

Alternation probability model: *y*_*t*_ = *d*_*t*_ for *t* = 2, …, *T* with *𝒪*_*AP*_ = {0, 1} and 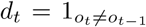

Transition probability model 1st Order: 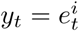 for *t* = 2, …, *T* with *𝒪*_*T P* 1_ = {0, 1, 2, 3} as 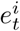 belongs to the set containing each possible transition from *o*_*t*−1_ = *i*.

Transition probability model 2nd Order: 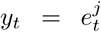 for *t* = 3, …, *T* with *𝒪*_*T P* 2_ = {0, 1, 2, 3, 4, 5, 6, 7} as 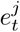 belongs to the set containing each possible transition from *o*_*t*−2_ = *j*.

### S2 Appendix. Variational inference algorithm

In this supplementary text we present the algorithm used to approximate log model evidence for subsequent Bayesian model comparison.

### A free-form variational inference algorithm for general linear models with spherical error covariance matrix

We consider variational inference for probabilistic models [58–60] of the form

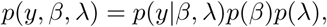

where

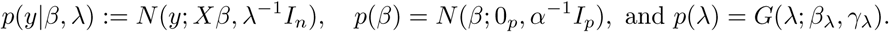

Here, *y* ∈ ℝ^*n*^ denotes the observed random variable modeling data, *β* ∈ ℝ^*p*^, *λ* > 0 denote unobserved random variables modeling regression weights and observation noise precisions, respectively, and *X* ∈ ℝ^*n*×*p*^ denotes a design matrix. The parameter-conditional distribution of *y* is specified in terms of a multivariate Gaussian density with expectation parameter *Xβ* ∈ ℝ^*n*^ and a spherical covariance matrix parameter *λ*^−1^*I*_*n*_. The marginal (or prior) distribution of *β* is specified in terms of a multivariate Gaussian density with zero expectation parameter 0_*p*_ ∈ ℝ^*p*^ and covariance matrix parameter *α*^−1^*I*_*p*_, where *α* > 0 denotes a precision parameter. Finally, the distribution of *λ* is specified in terms of a Gamma density in its shape and scale parameterization, where *β*_*λ*_, *γ*_*λ*_ > 0 denote the shape and scale parameters, respectively.

#### Model estimation

Application of the free-form variational inference theorem yields an algorithm that, upon convergence, furnishes an approximation to the data-conditional (posterior) parameter distribution of the form

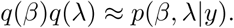

Here, the variational distributions take the form

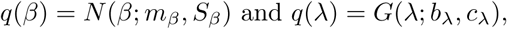

where *m*_*β*_ ∈ ℝ^*p*^ and *S*_*β*_ ∈ ℝ^*p*×*p*^ denote the converged variational expectation and covariance parameters, respectively, while *b*_*λ*_, *c*_*λ*_ > 0 denote the converged variational shape and scale parameters. Finally, the algorithm furnishes, upon convergence, the variational free energy lower bound

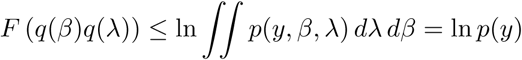

to the log marginal likelihood, also known as log model evidence. The algorithm takes the following form

##### Initialization

0. Set

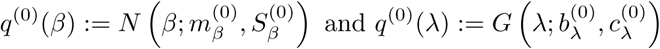

with variational parameters

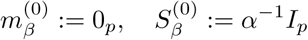

and

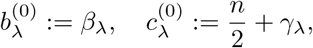

respectively. Define a convergence criterion *δ* > 0 and a maximum number of iterations *n*_*i*_.

##### Iterations

For *i* = 1, …, *n*_*i*_ or until convergence is reached

1. *<u>q</u>*(*<u>β</u>*<u>) update</u> Set

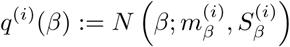

where

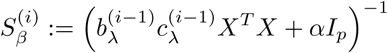

and

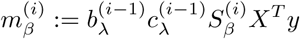
2. *<u>q</u>*(*<u>λ</u>*<u>) update</u> Set

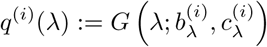

where

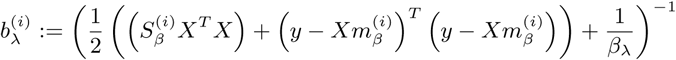

and

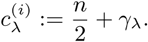

Note that 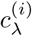 stays constant throughout.
3. *<u>F</u>*(*<u>q</u>*(*<u>β</u>*)*<u>q</u>*(*<u>λ</u>*<u>)) update</u> Set

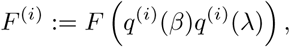

where

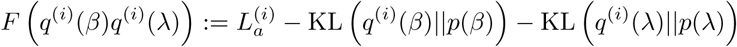

where with the digamma function 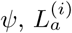 denotes the average likelihood term

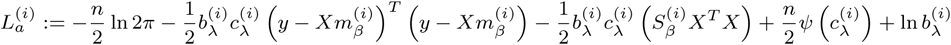

and KL(*q*(*x*)||*p*(*x*)) denotes the KL-divergence between the densities *q*(*x*) and *p*(*x*).
4. <u>Convergence assessment</u> If *i* > 1, evaluate

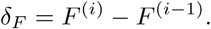

Then,

- if *δ*_*F*_ < 0, i.e., the variational free energy has decreased, issue a warning and end the algorithm,
- if 0 < *δ*_*F*_ < *δ*, i.e., the variational free energy has increased less than *δ*, end the algorithm and declare convergence,
- else go to 1.

#### Prior variational distributions

In order to select the probabilistic model of interest that minimizes Type II errors under the constraint of minimizing Type I errors the following test procedure was implemented. Data was simulated with low signal-to-noise levels (true, but unknown, *λ* = 0.001 and *β* = [1; 1]) and underwent z-score normalization, after which model retrieval was evaluated for a range of values for the precision parameter *α*. For data generated by the null model, a range of values of *α* was determined for which false positives were highly unlikely (exceedence probability *φ* = 1 in favour of the null model in every one of the 100 iterations). Next, for data generated by the non-null models, the value of *α* was selected which lied within the previously established range and for which the difference in log model evidence between null and non-null models was maximized. This procedure yielded the following prior distributions which were used in all described evaluations.

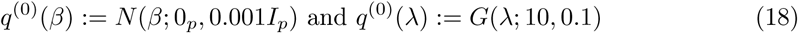

**S1 Fig.**
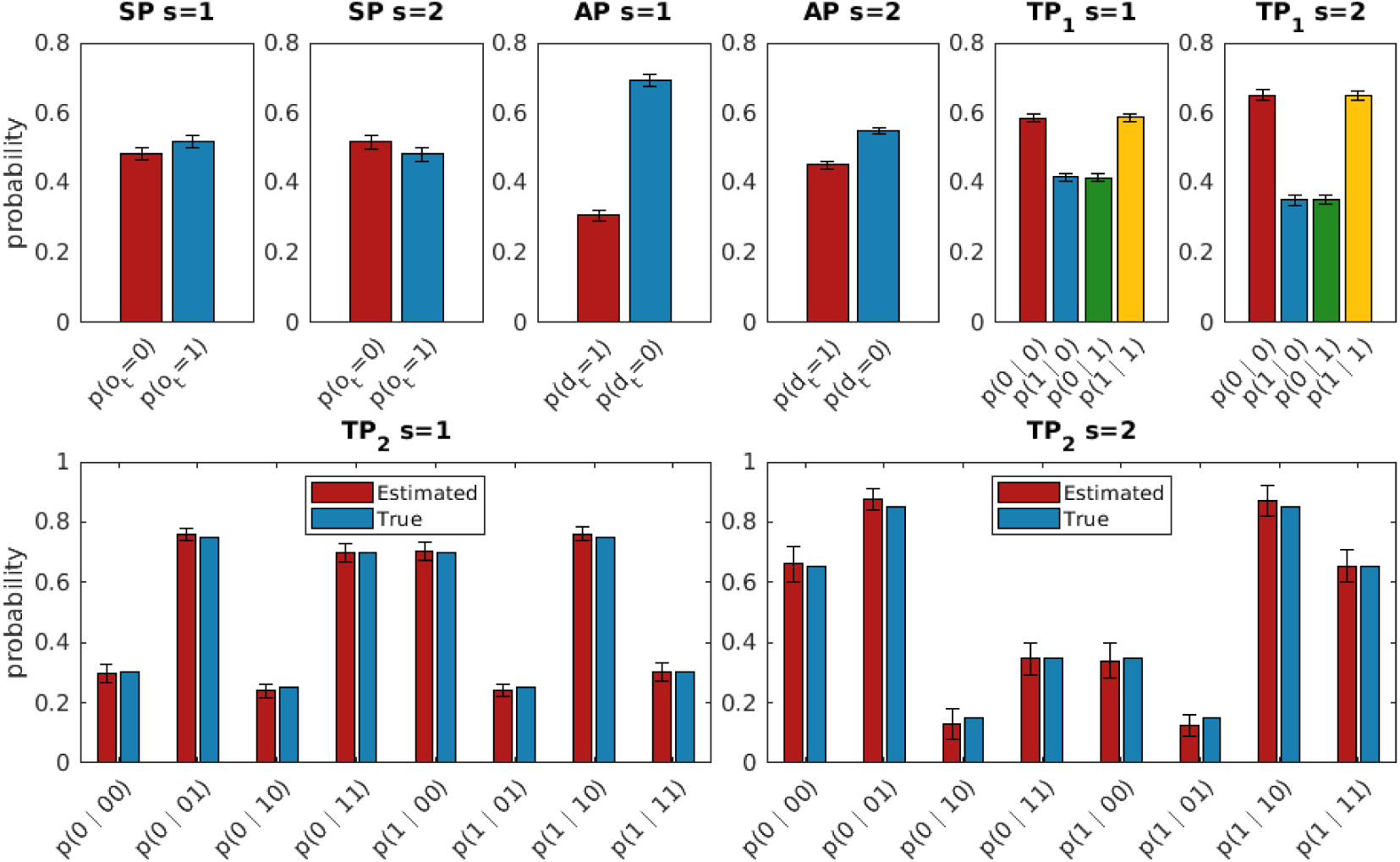
Average emission probabilities of the HMM. Top row: The average emission probabilities of the stimulus probability (SP), alternation probability (AP), and transition probability (TP) HMM for both states at the final timestep of each sequence. Bottom row: Comparison of the emission probabilities used for data generation and the average, normalized emission probabilities estimated by the HMM of type TP2. Error bars represent the standard error of the mean.

**S2 Fig.**
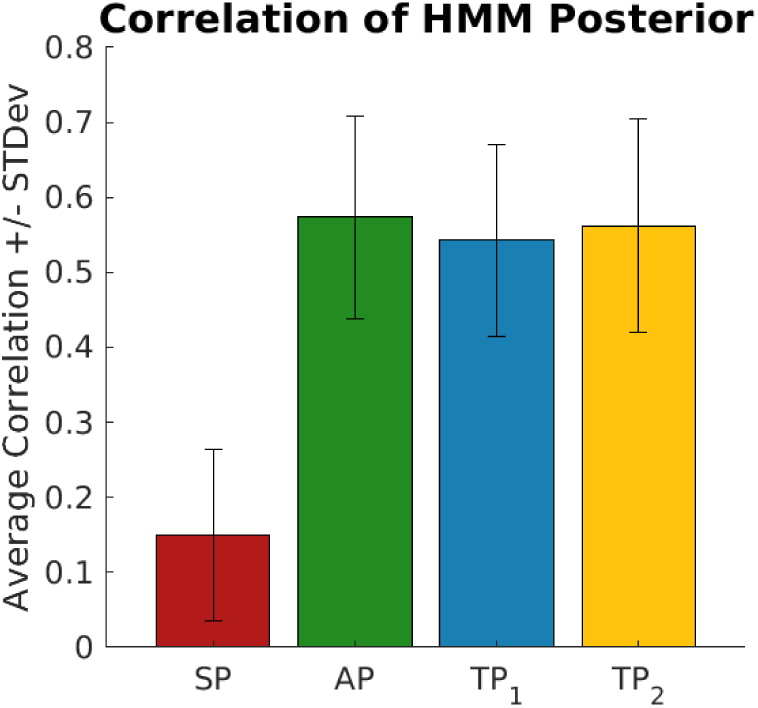
Average correlations between the HMM posterior and the regimes. Correlating the true regimes and filtering posterior over time confirms that AP and TP inference allow for the tracking of the fast and slow-switching regimes, while SP inference does not capture the necessary dependencies. SP: stimulus probability, AP: alternation probability, TP: transition probability

**S3 Fig.**
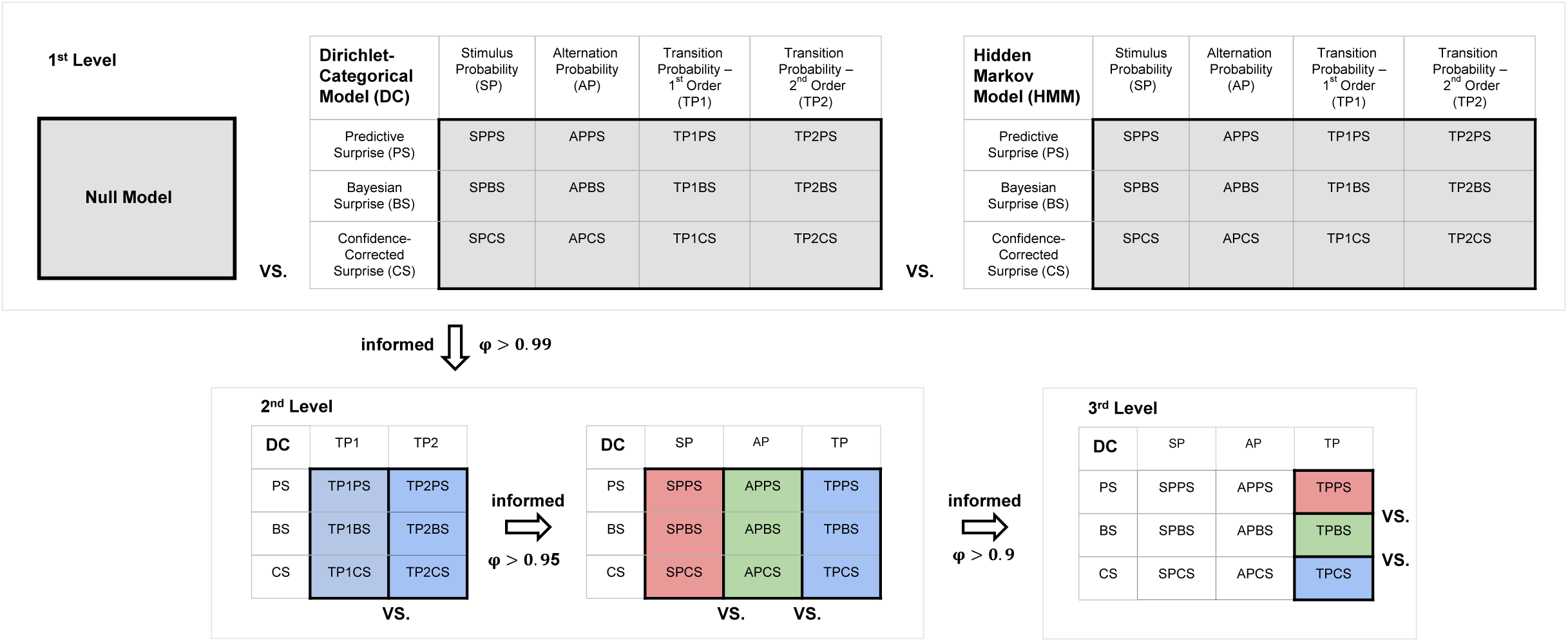
Schematic of the hierarchical random effects analyses. Hierarchical approach to familiywise Bayesian model selection. First level (depicted in the top row): The 12 DC models and the 12 HMM models were grouped into their corresponding model class family and compared via BMS against each other and an offset Null-Model. Second level (lower row, left rectangle): Within the DC model class, the two transition probability models TP_1_ and TP_2_ were grouped into families and the winner of the BMS was used for the comparison against the other two inference type models (Stimulus Probability (SP) and Alternation Probability (AP)). Third level (lower row, middle rectangle): The surprise readouts of the DC TP_1_ model were subjected to BMS and the resulting exceedence probabilities are reported in the main results.

